# WALLFLOWER, a polarized receptor-like kinase, alters cell wall properties impacting cell division and elongation in Arabidopsis

**DOI:** 10.1101/2025.07.15.664984

**Authors:** Patricio Pérez-Henríquez, Jessica N. Toth, Cecilia Rodriguez-Furlan, Rachel E. De La Torre, Dylan Wright, Haley Marks, Alexander A. Balandin, Jaimie M. Van Norman

## Abstract

Precise control of cell elongation and changes in directional growth are required for adaptive root responses. Anisotropic cell expansion in the root’s longitudinal axis must be coordinated among adjacent cell files and throughout the radial axis. However, links between activities across developmental axes remain obscure. We identified WALLFLOWER (WFL), a transmembrane receptor-like kinase that accumulates at the inner polar domain of epidermal cells in the root elongation zone. *wfl* mutants show deeper root waves and altered response to gravitropic re-orientation with increased cell length in specific root epidermal cell files. *wfl* phenotypes are rescued by WFL-GFP, but not by an intracellular truncation of WFL, which unexpectedly accumulates at the opposite (outer) polar domain. This suggests WFL function and its localization to the inner lateral plasma membrane domain are tightly coupled. Using advanced imaging techniques, including lifetime and atomic force microscopy, we found several mechano-chemical aspects of the *wfl* outer and inner cell walls are altered. This is consistent with a function for WFL in maintaining balanced mechanical properties between lateral cell walls during expansion. Overall, this work establishes a clear link between lateral protein polarization in peripheral tissues and longitudinal cell expansion required for adaptive organ growth.

## INTRODUCTION

Plant survival relies on adaptive growth and development, which occurs by continuously fine-tuning the rate and orientation of cell division and elongation triggering changes in growth direction in response to external conditions. In roots, the majority of longitudinal growth is driven by anisotropic cell expansion, which must be coordinated among cells in longitudinally adjacent files and also throughout cell types in the radial axis (Schindelman et al. 2001; Coen and Cosgrove 2023). Discordant elongation between neighboring cells could lead to shearing or rupture in extreme cases or less dramatic defects in cell and organ shape that may interfere with organ physiology and/or function (Bouton et al. 2002; Ubeda-Tomás et al. 2008; Hacham et al. 2011). Changes in directional growth (bending) are adaptive and intimately tied to cell expansion, as they require tight control of differential cell expansion in space and time. Adaptive growth ‘behaviors’ include hypocotyl elongation and unhooking in germination, changes in leaf angle and size, and root bending, among others. Our understanding of how the cellular processes required for adaptive growth are coordinated at the cell, tissue, and organ levels remains incomplete.

Adaptive root growth ‘behaviors’ are necessary for root function in uptake and anchorage and arise from differential regulation of cell division and elongation in response to a changing and heterogeneous soil environment (Robbins and Dinneny 2018; Wheeldon et al. 2022). In soil, roots often grow in a three-dimensional spiral pattern and can change growth direction or rate in response to obstacles, non-uniform nutrient or moisture availability, and/or the presence of biotic or abiotic stressors. When grown on the surface of an inclined substrate, roots display sinusoidal waving, that was initially proposed to result from the integration of gravitropic and thigmotropic (touch) responses (Okada and Shimura 1990; Oliva and Dunand 2007; Sedbrook and Kaloriti 2008). More recent theoretical frameworks suggest the importance of passive elasticity (Porat et al. 2024) and buckling of the elongation zone (Thompson and Holbrook 2004; Zhang et al. 2022). Despite these insights, the mechanisms underlying coordinated differential cell expansion are still largely unknown.

Precise spatiotemporal control of cell division and elongation result from refined molecular mechanisms (Bouchez et al. 2024; Yi et al. 2025) including a complex interplay between hormone, transcriptional, and cell polarity-based signaling. Cell polarity is defined as the asymmetric distribution of any cellular component, including protein accumulation to specific intracellular or membrane regions to exert local control on biological processes (Van Norman 2016; Ramalho et al. 2021; Jaillais et al. 2024). Polarized protein accumulation is required for many processes including: directional transport (Takano et al. 2010; Barberon et al. 2014; Janacek et al. 2024; Pérez-Henríquez et al. 2025), leaf and root patterning (Yoshida et al. 2019; Guo et al. 2021), polar cell growth (i.e. root hair formation)(Denninger et al. 2019), and tropic responses (Han et al. 2021). Proteins with polar accumulation may be soluble, membrane-associated, or membrane-spanning in nature. Leucine-rich repeat receptor-like kinases (LRR-RLKs) are transmembrane proteins broadly predicted to relay extracellular information into the cell to inform various developmental and immune processes (Bender and Zipfel 2023). A small number of polarly-localized LRR-RLKs have been identified and shown to function in root cell division and specialization (Alassimone et al. 2016; Campos et al. 2020; Rodriguez-Furlan et al. 2022; Goff et al. 2023; Chang et al. 2024; Rony et al. 2024).

The Arabidopsis root is an excellent system for studying cell polarity-informed developmental and growth events including cell proliferation, elongation, and specialization/maturation. These processes occur in the root’s longitudinal axis in defined regions termed the: meristematic, elongation, and differentiation zones (**Figure 1A**). In the transverse axis, root cells are organized in concentric rings around the central stele and are restricted in files of cells through the longitudinal axis. This organization allows for straightforward examination of cellular phenotypes and clear delineation of specific polarity domains within cells. In a longitudinal view, root cells display four broad polarity domains: shootward, rootward, inner, and outer (**Figure 1A**)(Dettmer and Friml 2011; Van Norman 2016; Ramalho et al. 2021; Jaillais et al. 2024). In transverse sections, the circumferential polar domain is also defined between adjacent cells in a ring (Ramalho et al. 2021). Spatial and directional information imparted by partitioned cellular components can direct cell division orientation and expansion during growth (Dong et al. 2023).

**Figure 1.**
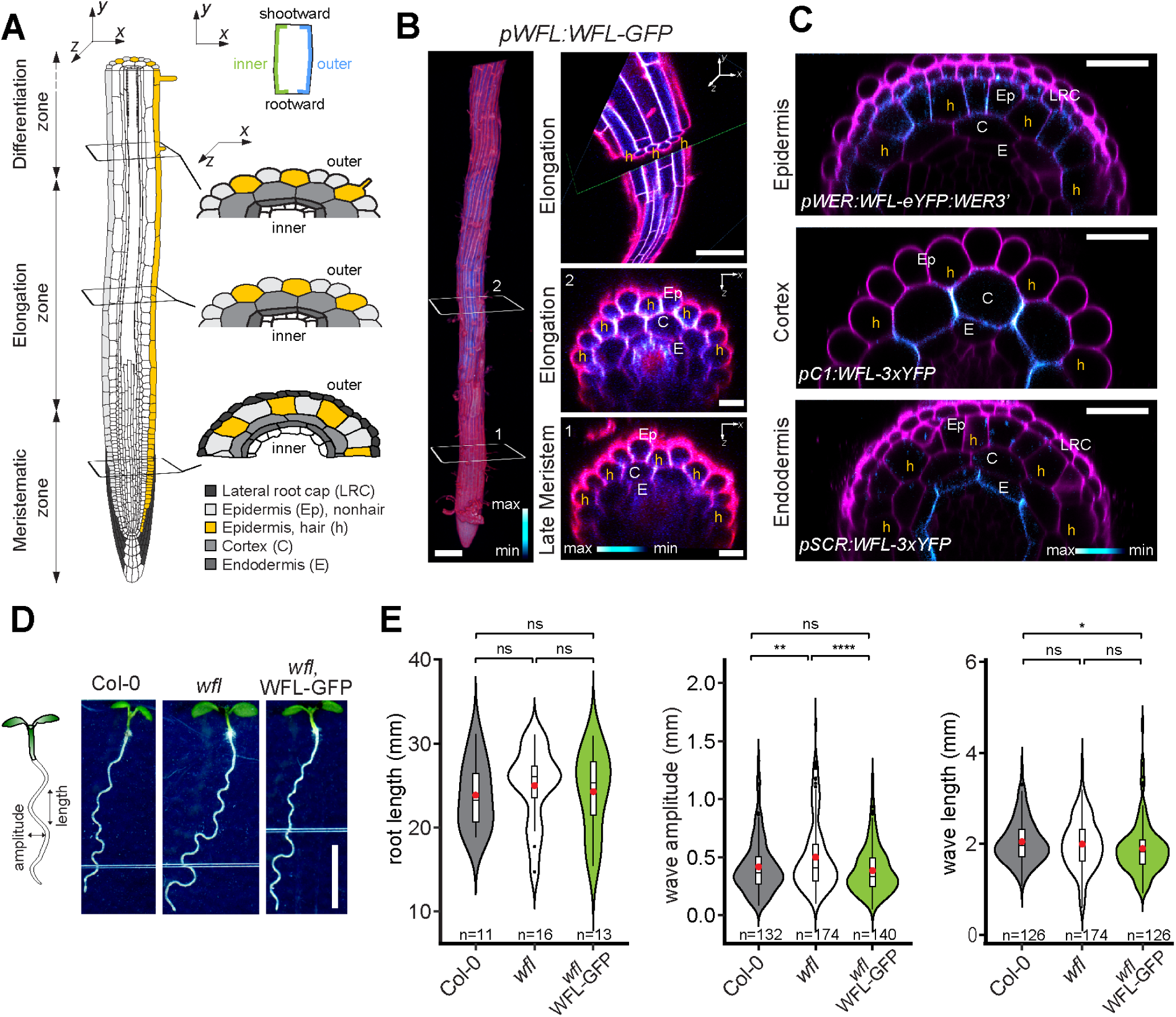
WFL accumulates at the inner polar domain in root cells and modulates root waving. **A)** Schematic representation of the Arabidopsis root tip with developmental zones and outermost cell types shown in (left) longitudinal and (right) transverse views. Epidermal hair (yellow) and non-hair (light gray) cells are indicated along with inner and outer (lateral) polar domains. (Upper schematic) Four plasma membrane polar domains represented in a single cell with inner/outer domains highlighted. **B)** Confocal images of a root expressing *pWFL::WFL-GFP* stained with propidium iodide (PI, magenta, GFP shown with intensity scale). *Left*, 3D projection shows higher accumulation of WFL-GFP in the elongation zone and numbered white boxes mark positions for *xz* images at right. Scale bar = 100 µm. *Right*, (top panel) intersecting *xz*-*xy* planes and (center and lower panels) *xz* planes show WFL-GFP in all epidermal cells in the elongation zone and primarily in hair cells in the meristematic zone. Scale bars = 100 µm (top) and 10 µm (center and lower) images. **C)** Confocal images of roots in *xz* planes stained with PI (magenta) and misexpressing WFL-GFP (intensity scale) under cell layer-specific promoters: *pWEREWOLF* (*pWER*) in the LRC and epidermis, *pSCARECROW* (*pSCR*) in the endodermis), *pCORTEX1* (*pC1*) in maturing cortex. Scale bars = 20 µm. Cell type abbreviations as indicated in A. **D)** *Left*, Schematic of root waving with wave amplitude and length indicated. *Right*, Representative images of 5 day-post-stratification (dps) Arabidopsis seedlings grown on plates at 20° inclination. Scale bar = 0.5 cm. **E)** Quantification of root length and wave features for a single representative replicate (of 3) for Col-0, *wfl*, and *wfl* WFL-GFP. Violin plots show data distribution with box plots indicating the first and third quartiles and dots showing outliers. ‘n’ indicates number of roots examined (left) and number of waves examined (center and right panels). The median and average values are shown by a line and red dot, respectively. Unpaired two-tailed *t*-test with Welch’s correction with ns = no significance (p-value > 0.05), * (p-value < 0.05), ** (p-value < 0.01), and **** (p-value < 0.0001). See also Figure S1 and Table S1.

Here, we identified a LRR-RLK that we named WALLFLOWER (WFL) due to its laterally polar accumulation in root cell types. Specifically, WFL accumulates at the inner polar domain of epidermal cells in the elongation zone and represses anisotropic elongation of non-hair epidermal cells in the longitudinal axis, impacting adaptive root growth behaviors, such as root waving and gravitropic re-orientation. Strikingly, removing the intracellular domains of WFL results in its accumulation at the opposite (outer) polar domain and does not complement *wfl* phenotypes as full length WFL does. This suggested WFL polar localization and activity are functionally linked. Using advanced imaging techniques in mechanobiology, including lifetime, atomic force, and brillouin microscopy, we find that the inner and outer walls of elongating epidermal cells maintain a balance in the mechanical properties of these walls, which is disrupted in *wfl* roots. We propose that WFL function at the inner polar domain is needed to preserve this balance during cell elongation. Overall, our work provides evidence of lateral cell polarity-based information in modulation of longitudinal cell elongation.

## RESULTS

### WFL-GFP is polarly localized in root epidermal cells

WFL is member of the RLK super family of plant receptor kinases and resides in subfamily LRRIII/VIIa along with other polarized LRR-RLKs, including KINASE ON THE INSIDE (KOIN, **Figure S1A**)(Man et al. 2020; Rodriguez-Furlan et al. 2022). To examine WFL accumulation, we performed confocal imaging on roots expressing WFL-GFP from the endogenous *WFL* promoter (*pWFL:WFL:GFP)*, which showed WFL-GFP accumulates strongly in the root elongation zone and weakly in the meristem (**Figure 1B, left and Figure S2B**). WFL-GFP accumulation is also detectable in meristematic epidermal hair cells and in the outermost layer of the lateral root cap (LRC). In the elongation zone, WFL-GFP accumulates most strongly in the pericycle and hair and non-hair epidermal cells and is weakly detected in the cortex (**Figure 1B** **and Figure S1C**). The observed WFL-GFP accumulation pattern is consistent with *WFL* transcript levels obtained from the single cell root atlas (**Figure S1B**) and aligns with our reporter for *WFL* promoter activity (*pWFL:erGFP*, **Figure S1C**).

In addition to accumulation in specific root developmental zones and cell types, WFL-GFP shows clear polar accumulation in outer cell types. In the lateral root cap, WFL-GFP accumulates to the inner polar domain of the plasma membrane (PM) (**Figure S1C**). In epidermal hair cells of the late (shootward) meristem, WFL-GFP preferentially accumulates at the circumferential polar domain (**Figure 1B**), while in the elongation zone, WFL-GFP accumulates at the circumferential and inner polar domains of all epidermal cell types (**Figure 1B**). Polar accumulation of WFL-GFP is also observed upon misexpression with cell layer-specific promoters. Specifically, WFL-GFP primarily accumulates at the circumferential polar domains in epidermal cells in the late meristem when expressed under the *WEREWOLF* promoter (*pWER)*, whereas it accumulates at the circumferential and inner polar domains when expressed in the endodermis *(pSCARECROW, pSCR)* and maturing cortex (*pCORTEX1, pC1*) (**Figure 1C**). This suggests that WFL polar accumulation doesn’t depend on cell identity and is informed, in part, by developmental stage. The spatiotemporal-specific accumulation of WFL suggests a link between its polar localization and function at different root developmental stages.

### *wfl* roots show altered root waving

To determine the function of WFL, we targeted *WFL* (At5g24100) for mutagenesis via CRISPR-Cas9 (Fauser et al. 2014). The *wfl-1* allele shows reduced *WFL* expression (**Figure S1D**) and is likely a null allele due to a premature stop codon early in the coding sequence (see methods section for details). We used information from the reporters to guide our phenotypic analysis with WFL-GFP in the pericycle prompting us to examine lateral root production in *wfl*, however, no change in lateral root density was observed (**Figure S1E**). Accumulation of WFL-GFP at the transition between meristem and elongation zones, and the shift in its accumulation, suggested WFL might function at this transition. When examining total root length, we detected a slight difference between *wfl* and wild type (Columbia, Col-0) roots, but that difference is not statistically significant (**Figure 1E, Figure S1E, Figure S3E)**. Additionally, examination of root length in a double mutant with the closest related LRR-RLK, named *WALLFLOWER-RELATED 1*, *WFLR1*, At5g53320, SALK_128780) showed that these related LRR-RLKs are not redundantly required for overall root growth (**Figure S1F-G**). With further examination of *wfl* roots, we noticed altered root waving compared to wild type (Col-0). This difference was exacerbated by growing seedlings on plates inclined 20°, which amplifies root waving. To characterize this, we quantified the most descriptive root wave features (Oliva and Dunand 2007), wave amplitude and wavelength (**Figure 1D-E**) and found that *wfl* root waves have a higher amplitude but similar wavelength. This indicates that *wfl* root waves are generated at the same frequency but are deeper, hinting at altered cell elongation and a transient or very local function for WFL in the root. This higher wave amplitude in *wfl* roots was not detectable in all growth conditions and was most obvious on inclined plates or on media that was firmer due to higher nutrient or agar concentrations (**Figure S2A**). However, on less firm plates without inclination, *wfl* roots showed higher curvature angles during gravitropic re-orientation (**Figure S1H)**. This suggests a loss of robustness in root growth behaviors in *wfl* in a range of growth conditions. Importantly, the abnormal *wfl* root growth phenotypes are rescued by endogenous expression of WFL-GFP (**Figure 1D-E** **and Figure S1H**). These phenotypes demonstrate that WFL function impacts aspects of root growth that are particularly important when changes occur in growth direction.

### WFL functions to repress cell elongation

Cell elongation is the primary driver of root growth and the epidermal growth control theory states that peripheral cell layers determine the elongation rate of an organ (Kutschera and Niklas 2007). The sinusoidal trajectory of root growth (waving) results from the modulation of cell elongation and buckling of the more elastic elongation zone (Thompson and Holbrook 2004; Zhang et al. 2022). Given that WFL-GFP strongly accumulates in the root elongation zone and not in more distal mature regions (**Figure S2B**), we hypothesized that changes in root waving in *wfl* are due to abnormalities in epidermal cell elongation. To test this, we performed *in situ* imaging to measure epidermal cell length while preserving positional information within root waves. After staining with propidium iodide (PI), multiple high magnification z-stack images were acquired, stitched together, and processed with MorphoGraph X (MGX) to project the surface signal into a 2D image (**Figure 2A**). This approach allowed us to distinguish curved peak regions from straighter interwave regions and hair from non-hair cells in Col-0, *wfl*, and *wfl pWFL:WFL:GFP* roots (**Figure 2A-B**). With these projections, we measured individual epidermal cell lengths in the longitudinal axis. Consistent with previous reports, we found wild type hair cells were slightly shorter in length than non-hair cells (Grierson et al. 2014). Unexpectedly, *wfl* and wild type hair cell length in peak and interwave regions were similar (**Figure 2C**), while non-hair cells were consistently longer in *wfl*. This longer non-hair cell phenotype was rescued by the endogenous expression of WFL-GFP (*wfl, pWFL:WFL:GFP*), suggesting WFL represses non-hair cell elongation.

**Figure 2.**
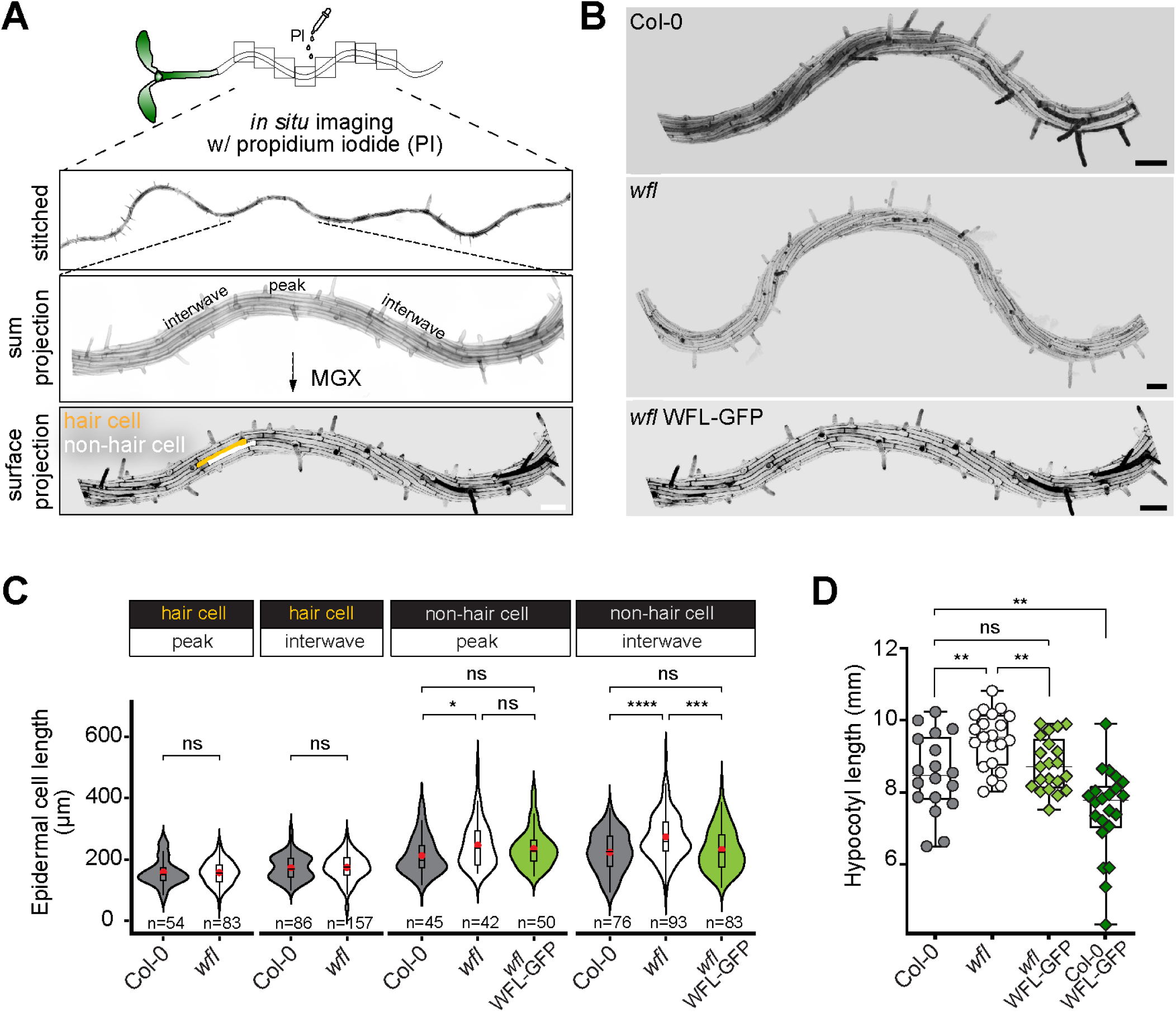
WFL represses elongation of non-hair epidermal cells. **A**) Schematic of *in situ* imaging approach for epidermal cell measurements within root waves. Roots were stained with propidium iodide (PI) *in situ* (without removing seedlings from the plate). Then, z-stack confocal images are obtained along the root’s length in tiles (open squares); then, stitched together to reconstruct a larger, continuous image area (upper root image). Root waves were divided into curved peak and straighter interwave regions. PI can be visualized by Sum (center root image) or Surface (lower root image) projections. In surface projections, obtained with MorphoGraphX (MGX), the PI fluorescence is projected onto a surface mesh, which allows clear delineation of epidermal hair (yellow) and non-hair (white) cells. Scale bar = 100 µm. **B**) Surface projections for *in situ* imaging of Col-0, *wfl*, and *wfl pWFL:WFL:GFP* roots. Scale bars = 100 µm. Seedlings were grown on inclined (20°) plates and imaged at 5 dps. **C**) Quantification of epidermal cell length in each genotype with number of cells indicated. Violin plots show data distribution with box plots indicating the first and third quartiles and median and average values indicated with a line and red dot, respectively. Unpaired two-tailed t-test. ns = no significance (p-value > 0.05), * (p-value < 0.05), and **** (p-value <0.0001). **D**) Quantification of hypocotyl length in Col-0, *wfl, wfl* WFL-GFP, and Col-0 WFL-GFP. Box plots show all data points and the box indicates the first and third quartiles, with the median value indicated as a horizontal line. Unpaired two-tailed t-test. ns = no significance (p-value > 0.05), ** (p-value < 0.01). See also Figure S2 and S3.

To examine WFL’s role in cell elongation more generally, we measured hypocotyl length in conditions, (sucrose-free growth media and dim light) that promote its growth based solely on cell elongation. In these conditions, *wfl* seedlings had longer hypocotyls than wild type (Col-0) and expression of WFL-GFP also rescued this phenotype (**Figure 2D**). Additionally, wild type seedlings expressing WFL-GFP (Col-0, *pWFL:WFL:GFP*, **Figure S2C**), had shorter hypocotyls than non-transgenic wild type (**Figure 2D**). This suggests repression of cell elongation through WFL is dose dependent. We detected WFL-GFP only at the rootward end of hypocotyls when expressed in *wfl* or in wild type (**Figure S3A-B**). We found no difference in *wfl* hypocotyl cell length, root growth rate or total root length compared to Col-0 on sucrose-free media under standard light conditions (**Figure S3C-E**). This indicates there is no difference in hypocotyl cell number or length due to changes in embryonic development, but instead that hypocotyl length differences arise during dim light-induced cell elongation. The hypocotyl length phenotypes when WFL is absent or overabundant are consistent with its role to repress cell elongation, particularly in conditions that induce more dramatic elongation.

Prompted by the hypocotyl phenotype in wild type expressing WFL-GFP, we closely examined the roots of these seedlings and found a slight root hair positioning defect. Root hairs typically initiate and grow out at a very defined distance above (shootward of) the hair cell’s rootward end (Denninger et al. 2019). However, wild type roots with additional *WFL* copies showed a higher frequency of root hairs initiating at a more rootward position, parallel with the cell’s rootward end (**Figure S2D-E**). No other root hair defects were observed and we could not detect any root hair positioning defect in *wfl* mutants or in *wfl* pWFL:WFL-GFP plants. To rule out the possibility that the downward shift in root hair position was due to some interference of GFP fused to WFL, we expressed WFL in Col-0 without a GFP tag and found a similar shift in root hair position (**Figure S2F**). Altogether, our findings on root epidermal cell length, hypocotyl length, and root hair positioning are consistent with a developmental context-specific role for WFL in modulating cell shape changes and elongation.

### WFL represses longitudinal anticlinal divisions of hair cells in the root meristem

In the root meristem, where proliferative division is the primary developmental activity, WFL localizes to the circumferential polar domain of epidermal hair cells. We previously showed that other LRR-RLKs in the LRRIII/VIIa subfamily, namely IRK and KOIN, function to repress cell division frequency and/or orientation in the root meristem (Campos et al. 2020; Rodriguez-Furlan et al. 2022; Rony et al. 2024), suggesting WFL may also have a role in cell division. To examine epidermal cell division in *wfl*, we performed 3D confocal imaging of root meristems (Campos and Van Norman 2021), which allowed us to assess cell division orientation and frequency (**Figure 3A**). In the *wfl* epidermis, we detected more frequent longitudinal anticlinal divisions (LAD), which result in bifurcation of one cell file into two. By examining intersecting *xy* and *xz* planes, we found that epidermal hair cells divide to produce additional non-hair cells, as the new cells contact just one underlying cortex cell (**Figure 3B**). The frequency of bifurcated epidermal cell files in *wfl* is restored to wild-type levels by expression of *pWFL:WFL:GFP* (**Figure 3B-D**). Increasing the number of non-hair cell files through hair cell division is a normal patterning strategy in Arabidopsis roots after germination (Berger et al. 1998). Consistently, ∼50% of wild type (Col-0) roots at 3 dps (days post stratification) display these epidermal LADs but we observe them in nearly 100% of *wfl* roots. Interestingly, a similar bifurcation of epidermal cell files was also observed in *wfl* hypocotyls at 3 dps, again at a higher frequency than observed in wild type (**Figure 3E-F**). This increase in epidermal cell number in the radial axis did not alter hypocotyl width (**Figure S3F**). Thus, in the absence of WFL, epidermal cells in the root and hypocotyl execute longitudinal anticlinal divisions more frequently than normal, again indicating a developmental-context specific role for WFL in root morphogenesis.

**Figure 3.**
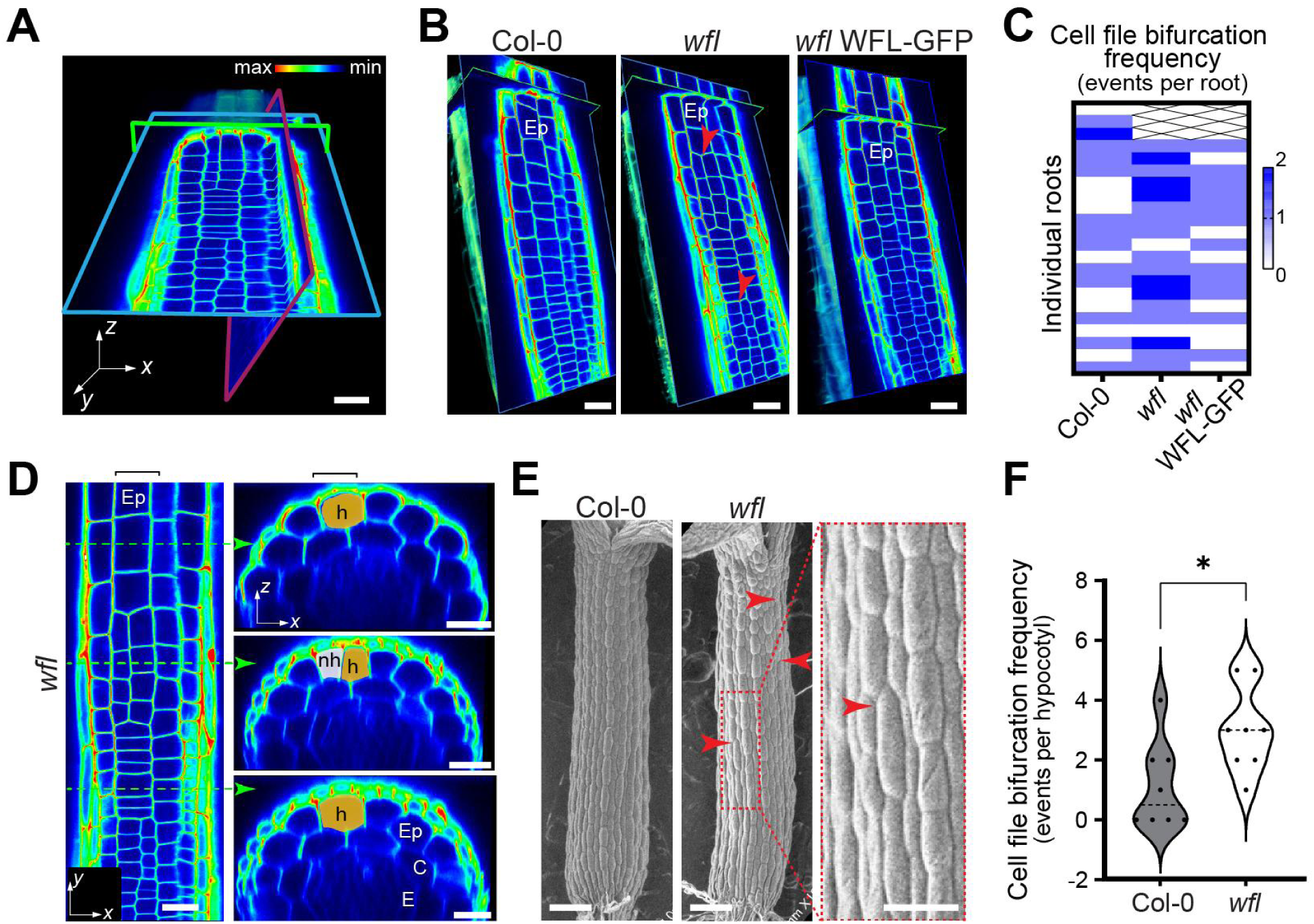
*wfl* roots and hypocotyls show additional divisions in epidermal cell hair cell files. **A, B)** 3D confocal images of PI stained roots (intensity color scale). **A)** Image displaying the clipped planes for views in *xy* (blue outline), *xz* (green outline), and *yz* (red outline). Scale bar = 20 µm. **B)** Representative images of epidermal (*xy*) planes of roots with longitudinal anticlinal divisions (LADs, red arrowheads) that bifurcate an epidermal (Ep) cell file to produce another (partial) cell file. Scale bar = 20 µm. **C)** Heatmap showing frequency of epidermal cell files bifurcation events in each genotype (n = 19-22 roots per genotype). Values from aggregated data in three independent experiments. Unpaired two-tailed *t*-test shows average frequency of these events in *wfl* is significantly higher (p-value < 0.01) than Col-0 or *wfl* WFL-GFP. **D)** Confocal image of *wfl* root epidermis (Ep) in the longitudinal axis (left), with green dotted arrows indicating the position of the transverse sections (right). Transverse sections depict a hair cell (h, yellow) indicated by contact with 2 underlying cortex (C) cells undergoing a LAD to produce a non-hair cell (nh, pink) that contacts only one underlying cortex cell. Black brackets indicate the epidermal cell file of interest in the *xy* (left) and *xz* (right) planes. Scale bars = 20 and 10 µm for *xy* and *xz* planes, respectively. **E)** Scanning electron microscope (SEM) images of hypocotyls at 3 dps in Col-0 and *wfl* with bifurcation events in epidermal cell files indicated (red arrowheads). Representative images (n = 8-9 hypocotyls per genotype). Scale bar = 100 µm for whole hypocotyls and 50 µm for zoomed-in image. **F)** Frequency of epidermal bifurcation events in hypocotyls (events per hypocotyl, n = 8-9 per genotype). Violin plot shows all data points with the median indicated by dashed line. Unpaired two-tailed *t*-test, * (p-value < 0.05). See also Figure S3.

### WFL-GFP is trafficked to plasma membrane by targeted exocytosis

Differences in WFL polar accumulation patterns in the root meristem and elongation zones suggests that WFL trafficking to the plasma membrane (PM) is dynamic and functionally important. Proteins with polar localization can be directed to the PM by targeted exocytosis and/or maintained at the PM by endocytosis and recycling (Rodriguez-Furlan et al. 2019; Raggi et al. 2020). To investigate how endomembrane trafficking contributes to WFL-GFP polar distribution, we examined its accumulation in roots following a series of chemical treatments. Brief treatment with Brefeldin A (BFA), an inhibitor of Golgi trafficking that affects exocytosis to the PM, generated intracellular agglomerations consistent with BFA bodies (**Figure 4A**). WFL-GFP accumulation in BFA bodies could result either from secretion of newly synthesized WFL-GFP or recycling of existing protein that has undergone endocytosis and is returning to the PM. To distinguish between these possibilities, we first treated roots with cycloheximide (CHX), an inhibitor of protein synthesis. After a 2 hours CHX treatment, WFL-GFP signal at the PM was markedly reduced, indicating that maintenance of WFL at the PM largely depends on continuous synthesis and secretion of new protein (**Figure 4B**). The strong decrease in WFL-GFP at the PM also suggests a high rate of protein turnover. If the existing pool of WFL-GFP at the PM is internalized and rapidly degraded, it would not be replenished when protein synthesis is blocked with CHX. To confirm this hypothesis, we pre-treated roots with CHX for 60 minutes then added BFA, and incubated for an additional 60 minutes. Under these co-treatment conditions, WFL-GFP accumulation in BFA bodies was almost completely abolished, with only a faint residual signal detected (**Figure 4A,B**). This indicates that most of the WFL-GFP detected in BFA bodies originates from newly synthesized protein entering the secretory pathway, rather than from recycling of existing protein.

**Figure 4.**
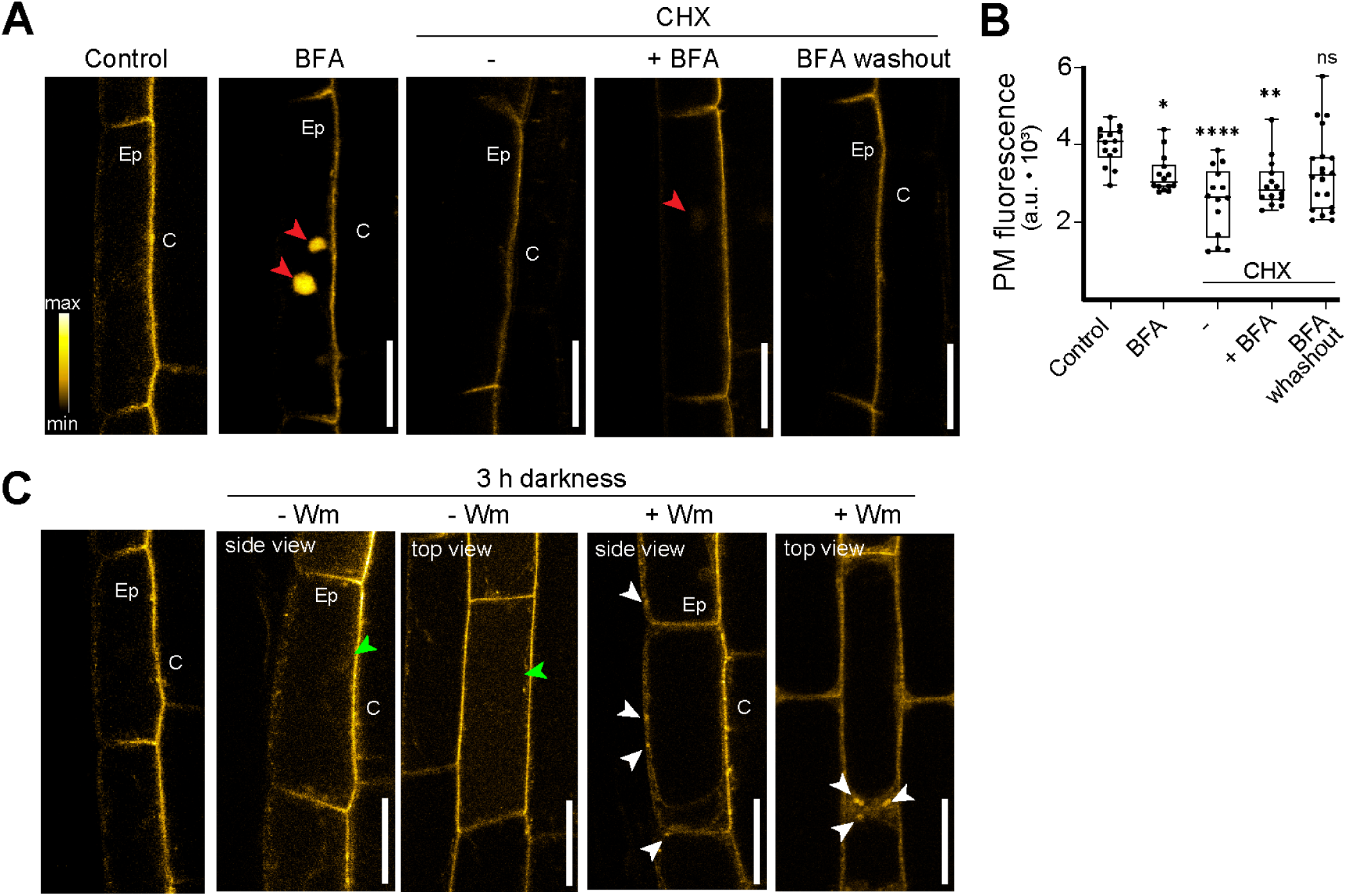
WFL is rapidly trafficked to and from the plasma membrane (PM). **A)** Confocal images of root epidermal (Ep) cells expressing WFL-GFP upon treatments: Control (solvent), Brefeldin A (BFA, 1h), Cycloheximide (CHX, 2h) and CHX co-treatments in the absence and presence of BFA (-/+BFA). BFA washout corresponds to the removal of BFA in the presence of CHX. Intracellular protein aggregates in BFA-bodies are denoted by red arrows. **B)** Quantification of GFP intensity at the PM after each treatment. Representative data shown from one of three independent replicates. The box plot shows all the data points with the box indicating the first and third quartiles and the horizontal line denotes the median. One-way ANOVA with Dunn’s multiple comparison test. ns= not significant (p > 0.05), * p < 0.05, ** p < 0.01, and **** p < 0.0001. **C)** Confocal images of root epidermal (Ep) cells expressing WFL-GFP upon treatment with solvent control (left panel, side view only) and 3h dark treatment in the absence (-Wm) or presence (+Wm) of Wortmanin (Wm). WFL-GFP accumulation at the vacuole (green arrowhead) or in the prevacuolar compartment (white arrowheads) with images shown in top and side view.

We also found that pretreatment with CHX dramatically reduced BFA-agglomerations and BFA wash out in the presence of CHX showed recovery of WFL-GFP signal at the PM suggesting that a proportion of WFL, however small, is recycling (**Figure 4A,B**). Because darkness induces internalization and trafficking of PM proteins to the vacuole but delays their luminal degradation allowing GFP detection at the vacuole lumen (Kleine-Vehn et al. 2008), we used a 3-hour dark treatment and detected increased WFL-GFP at the lumen in epidermal cells (**Figure 4C**). We next explored how WFL-GFP accumulation in darkness is impacted by Wortmannin (Wm), an inhibitor of phosphoinositide synthesis widely used to inhibit the endocytic trafficking of receptors from the PM towards the vacuole for degradation. A 2-hour Wm-treatment in darkness, revealed WFL-GFP in characteristic doughnut-shaped intracellular accumulations and considerable decrease in fluorescent signal at the vacuole lumen. Together, these results suggest WFL has a high turnover rate, being synthesized and degraded in a short time.

### WFL requires its kinase domain for its normal polarity and function

Like other LRR-RLKs with polar accumulation that we have identified, WFL is classified as an atypical kinase as it contains non-conserved residues in the catalytic domain (Castells and Casacuberta 2007). Nonetheless, the KOIN kinase domain is required for its function and polar accumulation at the PM, whereas the IRK kinase domain is largely dispensable for its polar accumulation and function (Campos et al. 2020; Rodriguez-Furlan et al. 2022). Therefore, we asked if WFL localization and/or function is affected by removal of its juxtamembrane (Jx) and kinase (K) domains (**Figure 5A**). In roots expressing *pWFL:WFLΔJxK:GFP*, we observed strong accumulation of WFLΔJxK-GFP in the elongation zone similar to full length WFL-GFP (**Figure 5B**). Strikingly, transverse optical sections revealed that WFLΔJxK-GFP is at the opposite lateral domain of epidermal cells - accumulating at the circumferential polar domain and the outer polar domain (**Figure 5B-C**). In the root tip, WFLΔJxK-GFP also accumulates at the outer polar domain of meristematic epidermal cells and cells of the LRC (**Figure S4A**). Furthermore, misexpression with cell layer-specific promoters showed that truncated WFL predominantly localizes to the circumferential and outer polar domain of many root cell types, except the endodermis, where it has nonpolar accumulation (**Figure S4B-C**). This is consistent with WFL polar accumulation being informed by an organ-level cue that generally orients WFL towards or away from the stele. More importantly, our data indicates that, while the WFL kinase domain isn’t strictly necessary for polar PM accumulation, it is needed to place WFL at the correct lateral (inner) polar domain.

**Figure 5.**
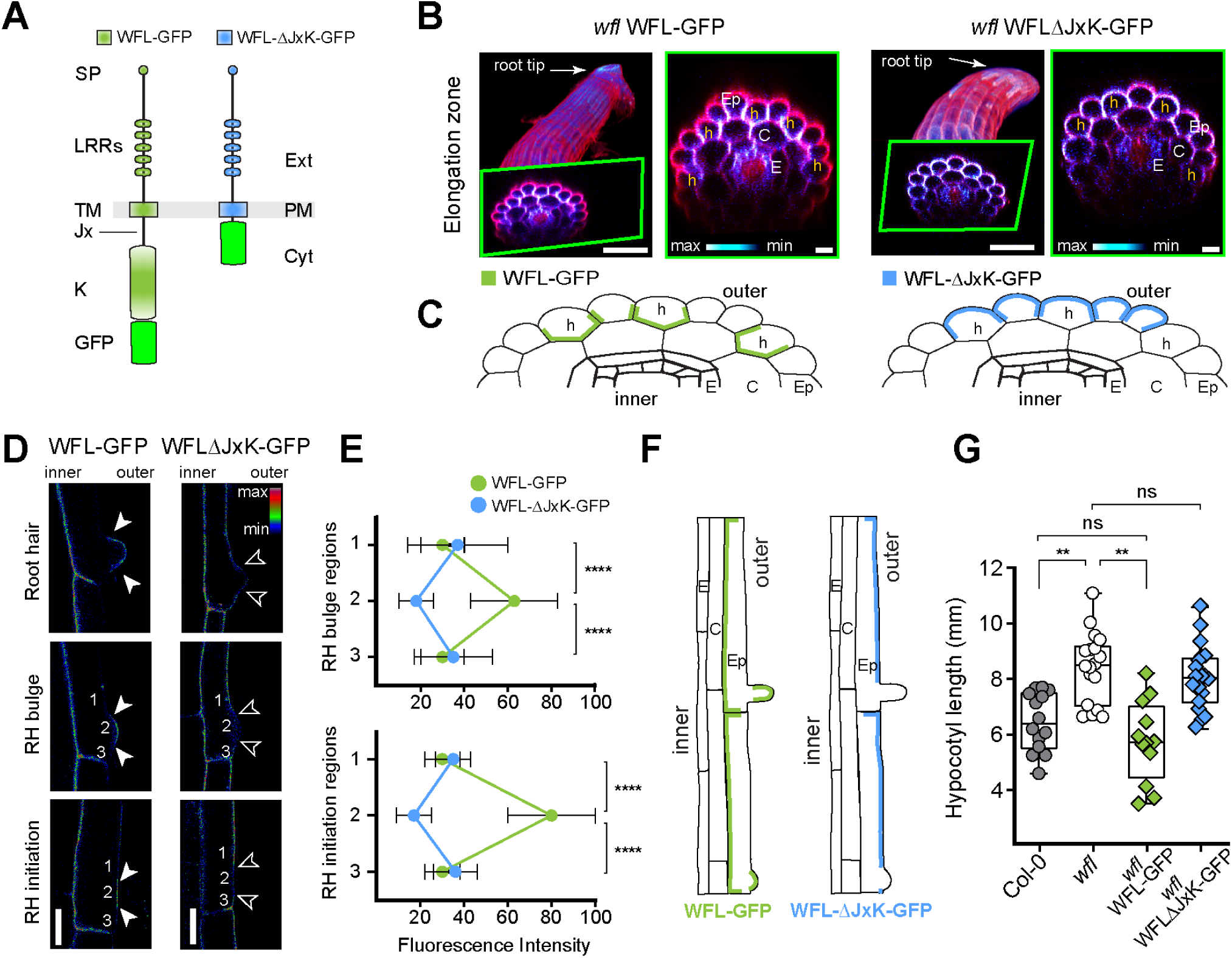
The WFL kinase domain is required for its localization and function. **A)** Schematics of protein domains for full length WFL (green) and truncated WFLΔJxK (blue). The transmembrane (TM) domain spans the plasma membrane (PM) separating the extracellular (Ext) signal peptide (SP) and leucine-rich repeat (LRR) domains from the cytoplasmic (Cyt) juxtamembrane (Jx) and kinase (K) domains. **B)** Confocal images of roots stained with propidium iodide (PI, magenta). (Left panels) 3D projections and (right panels) *xz* planes are shown for each genotype. Scale bars = 50 µm and 10 µm for left and right images, respectively. **C)** Schematic representations of WFL-GFP (green) and WFLΔJxK-GFP (blue) epidermal accumulation in the transverse root axis. Cell type abbreviations as indicated in Figure 1A. **D)** Confocal images during root hair (RH) formation of WFL-GFP and WFLΔJxK-GFP accumulation (in Col-0, shown with intensity color scale). Empty and filled arrowheads indicate the absence and presence of reporter accumulation, respectively. Numbers denote positions above (1), at the center (2), and below (3) RH initiation and bulge sites where fluorescence intensity was measured. Note that in the differentiation zone, WFLΔJxK-GFP is also detected in the outer polar domain of the cortex. Scale bar = 20 μm. **E)** Fluorescence intensity measured at the outer polar domain at positions indicated (**D**). Plots show mean values ± SD. t-test, **** p<0.0001. N = 15-20 roots per genotype, 2-3 cells per root. **F)** Schematic representations of WFL-GFP (green) and WFLΔJxK-GFP (blue) polar accumulation in root hair cells in the longitudinal axis. **G)** Quantification of hypocotyl elongation in various genotypes. Box plots show all the data points with the box indicating the first and third quartiles, and median value shown as a horizontal line. Unpaired two-tailed *t*-test. ns = no significance (p-value > 0.05), ** (p-value < 0.01). See also Figure S4.

While closely examining WFLΔJxK-GFP accumulation at the outer polar domain of epidermal cells, we noticed that it was conspicuously absent from sites of root hair formation. Given this and the downward shift in root hair positioning on plants with additional *WFL* copies, we compared the accumulation of WFL- and WFLΔJxK-GFP during root hair formation. We found WFL-GFP accumulates specifically at the center of root hair initiation sites and bulges, but not on the rootward or shootward sides of a root hair (**Figure 5D-E).** In contrast, WFLΔJxK-GFP is specifically diminished at the center of root hair initiation sites and bulges (**Figure 5F)**. This mirrored polarized accumulation in root hairs matches our observations of how these proteins accumulate at opposing lateral polar domains of cells in the elongation zone and is an unexpected consequence of WFL kinase domain deletion.

Finally, to determine whether WFLΔJxK-GFP was functional, we assayed hypocotyl elongation and found that, unlike full length WFL-GFP, it was unable to rescue *wfl* (**Figure 5G)**. Furthermore, WFLΔJxK-GFP was unable to rescue the root phenotypes that we observed in *wfl*, including the wave amplitude (**Figure S4D**) or curvature angle upon gravitropic reorientation (**Figure S1H)**. Altogether, these results indicate that the WFL kinase domain is not required for WFL polarized accumulation in general, but plays a crucial role in WFL function and localization at the inner polar domain.

### Asymmetry in lateral cell wall attributes is disrupted in wfl

How can we further link WFL function in cell elongation and its lateral accumulation? Abnormal *wfl* phenotypes together with the spatiotemporal polar accumulation patterns of WFL-GFP hinted at a connection to the cell wall. To explore this connection, we examined attributes of epidermal cell walls using several approaches. According to growth theory (Kutschera and Niklas 2007), cell wall acidification is a major factor in cell elongation and cell wall loosening. Using a ratiometric pH sensor, 8-hydroxypyrene-1,3,6-trisulfonic acid trisodium salt (HPTS)(Barbez et al. 2017), we assessed cell wall acidification in wild type root epidermal cells as they begin to elongate and observed a clear difference between the lateral epidermal walls, with outer walls having a lower pH (**Figure 6A, B**). This is consistent with accumulation of AHA1 (plasma membrane H(+)-ATPase 1) in the outer polar domain of these cells (**Figure S5A**). This indicates an asymmetry in the acidification of the outer and inner epidermal walls and suggests these lateral walls have different attributes. However, in *wfl* epidermal cells, the asymmetry in pH between lateral walls is not detectable (**Figure 6A, B**). This result suggests the lateral epidermal cell walls are more similar to each other in *wfl* than in wild type and is consistent with a direct link between localization and function of WFL at the inner polar domain.

**Figure 6.**
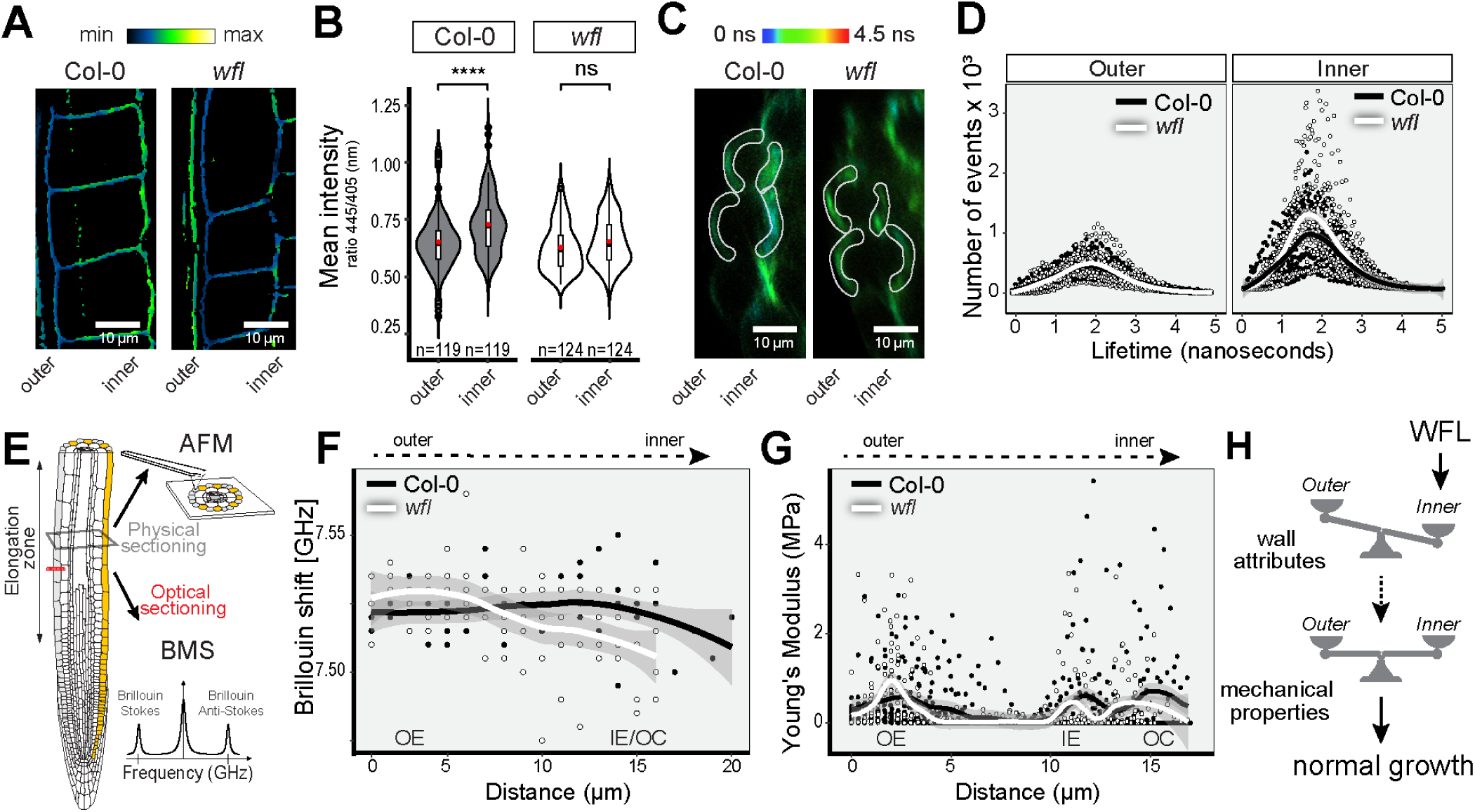
WFL functions to balance lateral cell wall properties in elongating epidermal cells. **A**) Ratiometric confocal images of root epidermal cells stained with the pH sensor HPTS (Barbez et al. 2017). Lower intensity (blue) represents lower pH. **B**) Quantification of HPTS ratiometric signal intensity. Violin plots show data distribution with box plots indicating the first and third quartiles and black dots indicate outliers. Median and average values are indicated with a line and red dot, respectively. Unpaired two-tailed *t*-test. ns = no significance (*p*-value > 0.05) and **** (*p*-value < 0.0001). n = number of cells, from 11 roots per genotype. C) Autofluorescence lifetime imaging (aFLIM) of root epidermal cells by optical cross-section. Regions of interest corresponding to the inner and outer walls (white outlines) were specifically examined. Blue coloration represents lower lifetimes (ns = nanoseconds). D) Histogram of apparent lifetimes obtained from FLIM images in outer and inner ROIs annotated in C. Continuous lines indicate data fitting to a Gaussian model. Dark gray area around each line indicates a 95% confidence interval. n = 50 cells per genotype collected from 10 roots per genotype. E) Root schematic displaying the physical sectioning for Atomic Force Microscopy (AFM) and optical sectioning with Brillouin–Mandelstam light scattering (BMS) microscopy. F) Brillouin shift values at different positions starting at the outer edge of the root. Continuous lines indicate a polynomial fitting. Dark gray area around each line indicates a 95% confidence interval. n = 5 roots per genotype. G) Young modulus values obtained by AFM. Values obtained from a line profile starting at the edge of the epidermal cell. Continuous lines indicate a polynomial fitting. Dark gray area around each line indicates a 95% confidence interval. n = 6 cells per genotype collected from 3 roots per genotype. H) Model for WFL function. Polar-localized WFL at the inner lateral side contributes to asymmetric (unbalanced) wall attributes; this leads to balanced mechanical properties between the outer and inner walls allowing for precise control of cell elongation in the longitudinal axis and normal root growth. Abbreviations: OE = outer epidermal wall, IE = inner epidermal wall, OC = outer cortex wall. See also Figure S5, Supplementary Video 1 and Supplementary Video 2.

pH asymmetry across epidermal cells in wild type implies structural and/or compositional differences exist between the lateral cell walls. Autofluorescence lifetime imaging (aFLIM) can be used to monitor cell wall components (Donaldson 2020) and dynamically detect their modifications (Escamez et al. 2021). With multi-photon and single-photon systems, aFLIM in wild type (Col-0) roots revealed lifetime differences in the lateral epidermal walls (**Figure 6C** and <u>Supplementary Video 1</u>, see methods section for details). The overall mean lifetimes in whole cross sections of *wfl* roots were lower than Col-0 (**Figure S5B**). More importantly, the local mean lifetimes between outer and inner sides of the epidermis were asymmetric in wild type but not in *wfl* (**Figure S5C**). Lifetime histograms at the inner epidermal wall vary significantly between Col-0 and *wfl* (**Figure 6D**). The pH and aFLIM data indirectly assay cell wall attributes and our results here imply structural and/or compositional asymmetries exist between the outer and inner epidermal walls in wild type roots and this asymmetry is disrupted in *wfl*.

### Balanced mechanical properties between lateral cell walls is disrupted in *wfl*

The above assays suggest that WFL at the inner polar domain functions to maintain asymmetrical attributes between lateral cell walls. However, in the elongation zone particularly, we envisioned that asymmetrical lateral cell wall properties would be problematic. During normal elongation, we reasoned that the lateral cell walls would need to have balanced mechanical properties to prevent shearing and rupture in extreme cases and generally to maintain cell and organ shape during growth. To directly examine properties of the epidermal walls, we probed mechanical properties of the lateral cell walls using two complementary approaches (**Figure 6E**). First, in optical sections, we scanned from the root edge inwards using Brillouin–Mandelstam light Scattering (BMS) microscopy (Kargar and Balandin 2021). BMS reports mechanical properties by measuring how light scatters after interacting with the sample; this generates a BMS frequency shift that is proportional to sample stiffness (Prevedel et al. 2019; Bacete et al. 2022; Zerin et al. 2025). At a whole root scale and consistent with previous findings, wild type roots showed higher BMS frequency shift (higher stiffness) in the stele than the outer layers (Bacete et al. 2022). In contrast, *wfl* roots displayed smaller BMS frequency shifts across the entire scanning area, indicating reduced stiffness (**Figure S5F-G**). Using a finer scanning resolution to focus on the epidermis, we observed no difference in BMS frequency shifts between the outer and inner sides of the wild type epidermis. This indicates symmetry (balance) in mechanical properties, namely stiffness, between the lateral epidermal walls. In contrast, the *wfl* epidermis displayed unequal BMS frequency shifts indicating a slightly stiffer outer wall and softer inner wall (**Figure 6F**).

We next examined the cell wall properties in the radial axis by probing physical sections of the root with Atomic Force Microscopy (AFM). Live roots were embedded in agar to maintain their structural integrity upon vibratome sectioning at the elongation zone (**Figure S5D and** <u>Supplementary Video 2</u>). Using AFM quantitative imaging mode, we obtained detailed maps of Young’s modulus values, a measure of stiffness, across the root and specifically across epidermal cells. Young’s modulus values across the root diameter displayed the same trend as the BMS data with increased stiffness towards the root’s center in wild type and *wfl* but the values at the center of *wfl* roots are substantially less (**Figure S5H**). With high-resolution probing across epidermal cells, we observed the outer and inner walls of wild type had the same stiffness, but in the *wfl* epidermis, the stiffness of the outer and inner walls is asymmetrical (**Figure 6G** **and Figure S5E**). Together, these direct assays of cell wall properties show that mechanical symmetry between the lateral epidermal walls is diminished in *wfl* with the inner wall appearing to be softer.

Taken together, our examination of cell wall features indicates that in wild type there are asymmetries in inner/outer cell wall attributes, but balance in their mechanical properties. In *wfl*, roots are overall softer, with attributes of the inner/outer wall being more symmetrical and mechanical properties unbalanced (**Figure 6H**). This suggests that, during cell elongation, WFL function is needed to maintain the balance between inner/outer wall mechanical properties and this function is intimately tied to its accumulation at the inner polar domain.

## DISCUSSION

Understanding how multicellular eukaryotes coordinate developmental processes across their bodies is a fundamental biological question. Root function requires effective soil exploration for plant productivity and survival necessitating adaptive root ‘growth behaviors’, including waving and bending, which are driven by differential cell expansion. In this context, coordination is typically envisioned as the extent to which cells elongate on different sides of the root. Because plant cells are bounded by a rigid cell wall, coordination between adjacent cells during differential growth is particularly important to maintain tissue and organ structure and function. But the wall surrounding a given cell is not necessarily uniform. In particular, and likely due to its protective function, the outer cell wall of the epidermis is substantially thicker than the inner wall (Dyson et al. 2014; Kelly-Bellow et al. 2023; Fino et al. 2025). However, this protective asymmetry presents a challenge: how to coordinatively balance the mechanical properties of the lateral (outer and inner) walls to prevent shearing and rupture. This suggests that activities at lateral walls must also be coordinated as cells expand longitudinally to ensure cell shape is maintained and link developmental events across orthogonal body axes. How coordination might be achieved at and between the cell, tissue, and organ levels in the longitudinal and radial axes is not clear.

In this work, we identified WFL, a transmembrane receptor kinase that is polarly localized in root cells. *wfl* mutants exhibit abnormal adaptive growth behaviors, including hypocotyl elongation and altered root waving and reorientation-induced bending, suggesting WFL functions to modulate cell elongation. In the root meristem, WFL localizes to the circumferential plasma membrane (PM) domain in epidermal hair cells and *wfl* mutants show early longitudinal anticlinal divisions in these cells. In the elongation zone, WFL accumulates to the inner polar domain in epidermal cells and represses non-hair cell elongation. Abnormal *wfl* phenotypes are restored by WFL-GFP and additional WFL copies in wild type have shorter hypocotyls and alterations in root hair positioning, suggesting WFL activity is dose-dependent. WFL arrives at the PM through targeted exocytosis and its trafficking is BFA-sensitive. Our results also indicate it is rapidly turned over and requires constant synthesis, consistent with a transmembrane protein whose activity is tightly regulated. An intracellular truncation of WFL (WFLΔJxK) fails to rescue the mutant and, unexpectedly, (mis)localizes to the opposite (outer) later polar domain, which links WFL polar accumulation to function at the inner side of cells. This link is further revealed by qualities of root epidermal cell walls. We observe altered epidermal cell wall mechanical properties in *wfl* leading us to propose that WFL acts to balance the intrinsically different qualities of the inner and outer walls during elongation (**Figure 6H**), acting locally at the inner side.

### WFL function and the balance of epidermal cell wall properties

Our observations with the pH sensor and cell wall autofluorescence lifetimes show that the biochemical attributes of epidermal lateral (outer/inner) walls are asymmetric and that this asymmetry is not maintained in *wfl*. At the same time, Brillouin and AFM microscopy indicate that *wfl* roots are generally softer and, at higher resolution, the lateral epidermal walls of wild type show a symmetry in mechanical properties that is not detected in *wfl.* To lengthen the root and maintain morphology, the lateral cell walls are loosened to allow for longitudinal expansion, while other walls do not loosen. To achieve uniform longitudinal expansion, symmetrical mechanical cell wall properties would be required and, if the lateral cell walls were identical, this would be straightforward - the cell could loosen outer and inner walls equally. However, the lateral epidermal cell walls are not the same; thus, we expect the outer and inner walls would be differentially modified/loosened to create symmetrical mechanical properties. Given that WFL accumulation peaks in the elongation zone and is localized to the inner PM domain in peripheral cell types, we propose that WFL functions in this specific place and time to fine-tune attributes of the inner wall to prevent it from loosening too much.

In general, WFL activity to locally fine-tuning cell wall stiffness can explain the different *wfl* phenotypes. This is straightforward for cell and organ elongation phenotypes. In the root meristem, increased longitudinal anticlinal divisions (LADs) in *wfl* hair cells occur, perhaps because without WFL at the circumferential polar domain, these cells are more susceptible to radial mechanical stress patterns (Hoermayer et al. 2024; Höfler et al. 2024). This could similarly explain early LADs observed in the *wfl* hypocotyl. However, repression of cell division activity by WFL would be less direct than we predict it is for IRK and KOIN, two related polarly localized receptor kinases (Campos et al. 2020; Rodriguez-Furlan et al. 2022; Rony et al. 2024). Moreover, WFL function to fine-tune cell wall stiffness could also explain the downward shift in root hair position observed in plants with expressing additional WFL copies. Root hair initiation requires substantial cell wall loosening of the cell wall at a very specific position in the outer epidermal cell wall (Vissenberg et al. 2001) and additional copies of WFL may impede this process, resulting in displacement of some root hair bulges.

### Mechanisms for WFL polar accumulation

The mechanism underlying WFL polarization within root cells is intriguing. WFL localization shows a dual polarity - being present at the inner PM domain of epidermal cells and at specific positions on the outer side during root hair initiation. This could be achieved through nonpolar trafficking of WFL to the entire PM and then selective removal. However, non-polar localization of WFL was not detected. We consistently detected polarly localized WFL in every developmental stage and upon treatment with BFA or Wortmannin, indicating polar localization of WFL is achieved by targeted exocytosis. This is similar to related LRR-RLKs that we have characterized, IRK and KOIN, which are also polarly localized by targeted exocytosis. However, IRK and KOIN polar localization is achieved through distinct pathways *i.e.* sensitive and insensitive to the golgi trafficking inhibitor BFA, respectively (Rodriguez-Furlan et al. 2022). Like IRK, WFL trafficking is also BFA-sensitive; therefore, they may share some commonalities in their polar accumulation mechanisms.

Further complexity arises due to the remarkable ‘mirror image’ accumulation of full length WFL-GFP and truncated WFLΔJxK. This contrasts with similarly truncated KOIN and IRK, where KOINΔJxK shows non-polar accumulation and IRKΔJxK displays correct polar accumulation (Rodriguez-Furlan et al. 2022). We concluded that a protein-intrinsic cue for polar accumulation is present or absent from the KOIN and IRK intracellular domains, respectively. However, WFLΔJxK presents an additional scenario and suggests we need to expand how we think about intra-molecular cues for polar LRR-RLK accumulation. In WFL, the information for polar (vs. nonpolar) accumulation may reside in its transmembrane or extracellular regions, while the cue to localize WFL to the correct (inner) polar domain resides in the intracellular region. Correct WFL localization may require interaction with specific PM environments (Pan et al. 2023) through the transmembrane and/or juxtamembrane domains, while WFL extracellular domains may also participate in polarization, via binding to specific extracellular or cell wall components, as proposed for other LRR-RLKs (Hématy et al. 2007; Feng et al. 2018). Observing mirrored polar accumulation of WFL and WFLΔJxK exposes considerable gaps in our understanding of protein polarization mechanisms, but also gives us the tools to investigate them in novel ways.

### Is WFL function determined by localization or intracellular domains?

WFL is categorized as an atypical kinase (Castells and Casacuberta 2007), presumed ‘kinase dead’, due to the presence of non-conserved residues at key positions in the catalytic domain, yet truncated WFL (WFLΔJxK) fails to rescue *wfl* phenotypes. This suggests the WFL intracellular domain is essential for its function, perhaps for downstream signaling. But as WFLΔJxK is also in the wrong place, it may not be able to function, perhaps due to a missing interaction partner or other functional constituent. Extra copies of WFL in wild type also resulted in mild abnormal phenotypes suggesting WFL activity is dose-responsive. These two observations suggest WFL’s presence at the right place and time are critical for its function, but don’t allow straightforward distinction between localization and functional domains. Indeed, they may not be mutually exclusive and determining the functional contribution(s) of WFL localization and/or the kinase domain separately requires a deeper investigation of WFL’s polarization mechanisms, its interaction partners, and downstream targets. Deciphering this can bring interesting insights to the ongoing discussion about polarization-dependent protein function (Nakamura and Grebe 2018; Muroyama and Bergmann 2019) in many aspects of plant growth and development. The precise molecular function of WFL in perception of informational cues from the apoplastic space and/or to aggregate direct cell wall modifiers will be part of our future work.

In summary, our work reveals a polarly-localized receptor kinase whose function is needed to balance mechanical properties of lateral epidermal cell walls to fine-tune cell elongation, ultimately, impacting organ-level adaptive growth. This work demonstrates an important link between lateral cell polarity-based information and modulation of longitudinal cell expansion.

## MATERIALS & METHODS

### Plant materials and growth conditions

The *Arabidopsis thaliana* Columbia-0 (Col-0) ecotype was used as the wild type. All seedlings were grown on plates in a Percival growth chamber at 22°C under a 16 h light/ 8 h dark photoperiod with 60-110 µmol/m^-2^s^-1^ of light intensity with details indicated for specific experiments. In vitro growth was carried out in 1X Murashige and Skoog (MS) basal salts (Caisson Lab®, Cat. Num. MSP01), 0.05% MES, 1% sucrose, pH 5.7 and 1% Agar (Difco Agar, BD®, Cat. Num. 214530), unless otherwise noted. Seeds were surface sterilized with chlorine gas and stratified on plates at 4°C for 2-3 overnights and then put into the growth chamber. Plants were propagated in a growth room either with 24 h light or 16 h light/ 8 h dark photoperiod at 20-24°C.

### Vector Construction and Plant Transformation

Transcriptional (promoter activity) and translational reporters were constructed by standard molecular biology methods and utilizing Invitrogen Multisite GatewayⓇ technology (Carlsbad, USA). A region 4.1 kb upstream of the *WFL* (At5g24100) start codon was amplified from Col-0 genomic DNA and recombined into the Invitrogen pENTR^TM^ 5’-TOPO(Ⓡ TA vector. For *WFL* promoter activity, the promoter drove endoplasmic reticulum-localized green fluorescent protein (erGFP) as previously described (Van Norman et al. 2014). For translational fusions, the genomic fragment encoding *WFL* from the ATG up to, but excluding the stop codon (including introns, 2.0 kb), was amplified from Col-0 genomic DNA and recombined into the Invitrogen pENTR^TM^ DIRECTIONAL TOPOⓇ (pENTR-D-TOPO) vector and fused to a C-terminal GFP tag (unless otherwise noted) as previously described (Van Norman et al. 2014). Specific primers for *WFL* cloning are listed in Table S1. WFL-GFP and WFLΔJxK-GFP were driven by cell type-specific promoters (*pSCR_2.0_, pCO_2_, pC1, pUBQ10,* and *pWER)* as previously described (Lee et al. 2006; Campos et al. 2020). Due to the relatively low fluorescent signal of *pWFL:WFLΔJxK-GFP,* misexpression reporters for these truncations were reconstructed with 3xYFP. *pC1* was received in (Gateway Compatible) pENTR^TM^ P4P1R TA vector from the lab of Philip Benfey, Duke University (Durham, NC, USA). The epidermal misexpressed fusion protein, *pWER:WFL-eYFP:WER3’,* was generated as previously described (Campos et al. 2020).

The various Gateway compatible fragments were recombined together with the dpGreen-BarT or dpGreen-NorfT (Norflurazon (Norf) resistance) destination vector (Lee et al. 2006). The dpGreen-NorfT vector was generated by combining the backbone of dpGreen-BarT with the p35S::tpCRT1 and terminator insert from pGII0125. Within the target region of the dpGreen-BarT, one AclI site was mutated with the QuickChangeXL kit (Stratagene). Plasmids were amplified in ccdB-resistant *E. coli* and plasmids prepped with a Bio Basic Plasmid DNA Miniprep kit. 34 uL of the modified dpGreenBarT and unmodified pGII0125 were digested with 1 µL each FspI and AclI in CutSmart buffer (NEB) for 1hr at 37°C. Digests were subjected to gel electrophoresis on a 1% agarose gel. The 5866 bp fragment from the dpGreenBarT and 2592 bp fragment from the pGlIII0125 were extracted with a Qiagen MinElute Gel Extraction kit. The fragments were then ligated at 1:1 volumetric ratio (20ng vector; 8.8ng insert) using T4 DNA ligase incubated at 16°C overnight before transformation into ccdB-resistant *E. coli*. Expression constructs were then transformed into Col-0 plants by the floral dip method(Clough and Bent 1998) using Agrobacterium strain GV3101 (Koncz et al. 1992) and transformants were identified using standard methods.

For each transgenic reporter line, up to 15 independent T1s were isolated and propagated. T2 lines with a 3:1 ratio of resistant:sensitive seedlings, indicating the transgene is inherited as a single locus, were allowed to self (3-5 lines minimum) and among the subsequent T3 progeny, those with 100% resistant seedlings, indicating that the transgene was homozygous, were used in further analyses. For each reporter, at least three independent lines with the same relative expression levels and localization pattern were selected for imaging by confocal microscopy.

### CRISPR-induced mutagenesis

A candidate T-DNA insertional allele of *WFL* was obtained from the ABRC (Arabidopsis Resource Center), SAIL_1170_A12, but was discarded as no heterozygous or homozygous mutant individuals could be identified in multiple batches of seeds. An allele (*wfl-1*) was generated using CRISPR-Cas9 technology and used for all phenotypic analyses. CRISPR-induced mutagenesis was performed as described in (Fauser et al. 2014), a single guide RNA (5’-TTTAACGGTAGTATTCCCGCGGG) was selected in exon 2 of *WFL.* Single locus heritability was tested with 3:1 ratio of resistant:sensitive seedlings segregation in the T2 population. These plants were subsequently examined for lesions in *WFL* in proximity to the guideRNA binding site. In two individual transgenic lines, single point mutations were obtained at the same position. A single T (*wfl-1)* or a single A (*wfl-2*) was inserted 486 nucleotides after the start codon, which results in 4 novel amino acids and a premature stop codon. Because these alleles have the same consequences on the protein coding region, generating a stop codon prior to an intron and the coding region for the transmembrane domain, it is unlikely that any partially functional protein could be produced; thus, we focused our analyses on *wfl-1*.

### Confocal microscopy

All confocal images were acquired with laser scanning confocal microscopes using either a Leica SP8 (housed in the Van Norman lab at University of California, Riverside, UCR) or a Zeiss 980 (housed in the Van Norman lab at UC, Los Angeles, UCLA) with water immersion, 40X objective, NA = 1.2. Fluorescent signals were captured using the following settings: propidium iodide (PI; excitation 543/561 nm, emission 580-660 nm), GFP (excitation 488 nm, emission 492-530 nm), and YFP (excitation 514 nm, emission 515-550).

### 2D Confocal image processing

Post-acquisition image processing was performed with proprietary software Zeiss ZEN® 3.7 software, Leica Application Suite® X (LASX) software, or with the open source Fiji (Fiji is Just ImageJ) v1.54. Final multi-panel figures were assembled using Adobe® Illustrator and Adobe® Photoshop. Plasma membrane fluorescence intensity in chemical treatments were processed using ImageJ. The intensity was corrected by background readings using the formula: Integrated Density - (Area of selected cell x mean values of background readings). Plasma membrane fluorescence intensity in root hairs was obtained using LASX. Position 1, 2, 3 corresponds to the area above, center or below the root hair initiation domain or bulge. Measurements were performed in 2-3 cells per root in a total of 15-20 roots per replicate.

### 2.5 and 3D confocal image processing

All tiled images were acquired with confocal microscope Zeiss 980 (UCLA) and processed at a high-performance PC running Windows 10 Enterprise equipped with a CPU AMD Ryzen Pro 3955WX with 16-cores 3.9GHz and GPU Nvidia RT A5000. Tiled images were stitched together before further processing with the Zen desk v3.8 software using the processing function ‘Stitching’ and following parameters. FUSE TILES: Yes, DIMENSION REFERENCES: All by reference (pick a z-position with clear signal), EDGE DETECTOR: Yes, MINIMAL OVERLAP: 5%, MAXIMAL SHIFT: 5%, COMPARER: Best, GLOBAL OPTIMIZER: Best.

Stitched images were then processed for single-channel 2.5D projections (surface projections) or dual-channel 3D projections. <u>2.5D projections or surface projections</u> were calculated with MorphoGraphX v2.0. For this, 16-bit images were acquired with a stepsize of 1 µm. Lower step size dramatically increased the file size without improving the epidermal signal projected to the surface. The analysis assisted by MGX was the following: First, images were converted from CZI to TIFF using Fiji. Then, a Gaussian blur filter was used [XYZ sigma = 0.3-1.2 µm]. Voxels were edited to erase the blurry noise around the image. Edge detection was run with THRESHOLD=default (10000). Then, Mesh was created with the marching cubes surface process using CUBE SIZE = 5. Mesh was manually trimmed eliminating mesh built at the base of the z-stack to avoid unnecessary processing in the next steps. Importantly, mesh processing was two rounds of subdividing and smoothing the mesh.

Finally, signal was projected onto the surface with MIN DIS = 3 µm and MAX DIST = 8 µm. *<u>3D</u> <u>projections</u>* were performed with the Arivis Vision4D v 4.1.1 software using 8-bit images acquired with a step size of 0.3 µm. GFP and PI channels were visualized with Rainbow and red LUT. Images were processed in Arivis with the following workflow: 3D RENDERING: Transparency, 4D OPACITY SETTINGS: Opacity curves = Transparency. COLOR SETTINGS: Gamma = 1, and 4D CLIPPING: plane *xz* = Invisible or Textured/Opaque.

### FLIM confocal microscopy

Autofluorescence lifetime imaging (aFLIM) was performed in 2 different systems. For z-stack images (<u>Supplementary Video 1</u>), a multi-photon Leica SP8 FALCON DIVE (40X IRAPO Water NA=1.1) with a 80MHz 1040nm pulse laser was used. For optical cross-section images, a confocal Leica SP8 FALCON FLIM (40X APO CS2 Oil NA=1.1) with a 80MHz pulsed white light laser tuned to 488nm was used for excitation. Broadband emission was collected using acousto-optical beamsplitters (AOBS) set from 550-800nm for the multiphoton nDD and 500-780nm for the HyD TCSPC detectors. All images were processed with Leica FALCON FLIM software . All images were initially explored with phasor analysis to establish a representative gradient of lifetimes (Digman et al. 2008). For quantification of the optical cross-section images, histograms of autofluorescence lifetime (in nanoseconds) are obtained from the whole image and from specific regions of interest at the outer and inner side of the epidermis, as shown in Figure 5C. Histogram data is fitted with a Gaussian curve model directly from the raw data, matching results obtained with a LOESS polynomial fitting model, where mean lifetimes are calculated as the weighted average from histogram data.

### Brillouin microscopy

Brillouin spectra (Kargar and Balandin 2021) were obtained using a JRS Table Stable TFP-2HC high contrast interferometer. The measurements were performed with a 4mm mirror spacing giving a free spectral range of ∼70 GHz and a spectral resolution of ∼30 MHz. The samples are mounted on a micropositioning stage, which enables precise lateral adjustments over the sample. A 532-nm, 2 W Coherent Verdi V-2 laser was attenuated to 1.6 mW and focused onto the sample using a 10X long working distance objective lens (NA=0.28). Seedlings were vertically mounted on a coverslip and covered with 2mm of the same semisolid agar media used for plant growth. The laser was directed to the outer edge of the root elongation zone, located 250 µm shootwards from the root tip. Each spectrum was accumulated over 30 seconds. For half-root measurements, the laser was repositioned inwards with a step size of 5 µm. For specific measurements within the epidermis layer, a step size of 1 µm was used. Spectral data were fitted with a Lorentzian function to determine the Brillouin frequency shift (BMS frequency shift). While we chose to report the BMS shift, identical results were obtained when calculating the Brillouin Elastic Contrast, as previously defined (Bacete et al. 2022).

### Atomic Force Microscopy

AFM microscopy was performed on the NanoWizard® 4a BioScience (Bruker, Inc.), mounted on an optical macroscope and HybridStage™ with a z-range of 200 µm. All images were acquired in liquid using the Advanced Quantitative Imaging™ mode. Samples were prepared with 5 dps (days post stratification) roots embedded in 7% low gelling temperature agarose (Sigma-Aldrich, Cat. Num. A-4018). Physical root cross-sections were obtained with Vibratome (Microm HM650V) using the following settings thickness/trim = 200 µm, frequency = 75 Hz and amplitude = 1 mm. For entire cross-section imaging, AFM imaging settings were: setpoint = 1 nN, z-step = 7000 nm, z-speed = 200 µm/s, and using the cantilever PFQNM-LC-V2 (tip radius = 70 nm, Bruker, Inc.) For imaging the epidermal cells, the settings were: setpoint = 1 nN, z-step = 2000 nm, z-speed = 75 µm/s and using the PPP-CONT (tip radius < 10 nm, NANOSENSORS™). Young’s modulus maps were obtained by analyzing the force-curves with the software ‘JPK Data Processing’ v6.4.35 in batch processing mode using the Hertz model (Tip shape = Paraboloid for PFQNM-LC-V2 and Tip shape= Triangular Pyramid for PPP-CONT cantilevers). The resulting Young’s modulus map was later processed with Gwyddion v2.69 (https://gwyddion.net). Line profiles (thickness=3 pixels) were drawn across the entire root diameter and across the epidermal cell width. Data from line profiles was then exported and processed in RStudio (*ggplot2* package) for graph building. A local regression fitting model (LOESS) was used to obtain the trend curves.

### Scanning electron microscopy

Hypocotyls from seedlings at 3 days post-stratification (dps), grown in normal light conditions, were imaged using the Hitachi TM4000 Plus II with an electron gun voltage of 15 kV (at UCR). Whole seedlings were displayed directly on the mounting platform on top of double-sided tape. The sample was flattened down by carefully tapping down the root and the cotyledons. Images were obtained at 100X magnification in mixed detection mode BSE/SE which combines backscattered electron imaging (BSE) and secondary electron (SE) detection. Default vacuum conditions were used as they are specially designed for hydrated samples.

### Root phenotypic analyses

Seedlings are grown for 5 dps at 20° inclination to enhance root waving as indicated by Tan *et al* (Tan et al. 2015). Root wave features, as described by Oliva *et al* (Oliva and Dunand 2007), were measured in Fiji from images captured on a flatbed scanner. Plates were positioned at 20°C inclination at day 0 in the growth chamber for seedling germination and growth. Root length was obtained by tracing the root with the segmented line tool in ImageJ from images of the plates. Lateral root density was obtained from seedlings grown vertically through the agar in plates and was calculated as the number of emerged lateral roots divided by primary root length. Root hair bulge position measurements were performed on seeds at 4 dps grown on 0.5 X MS media following published protocols (Masucci and Schiefelbein 1994). Root hair bulges were binned into 2 categories with the following parameters: “normal” if there was a measurable distance from the emerging root hair to the rootward edge of the harboring epidermal cell and “rootward shifted” if there was no measurable distance. Root curvature angle was obtained by reorienting root growth in 180° degrees. Seedlings were grown on the surface of 0.2X MS in plates with 0° of inclination for 5dps and grown for another 24h after re-orientation. These growing conditions favor straight root growth. Straight roots prevent root waving affecting the curvature angle formed during gravitropic reorientation. The curvature angle was obtained with the Fiji plug-in claw curvature FeducciA placing two points (A,B), each aproximately 2 mm away from the minimum point of the curvature (X).

### Hypocotyl phenotypic analyses

Hypocotyl length assays were performed on seedlings growth on media without sucrose. Sucrose is omitted to avoid interference with growth rates and de-etiolation response. For dim light response assays, seeds were first germinated in light at 60 µmol/m^-2^s^-1^ for 2 days, then exposed to dim light (5 µmol m^-2^ s^-1^) for 4 days. Plates were positioned horizontally throughout the experiment. After dim light treatment, hypocotyls are transferred to and aligned on a humid transparent plastic sheet to obtain images (acquired on a flatbed scanner). Light intensity was measured with a LI-250 light meter (LI-COR ®) equipped with a quantum sensor. A dim light environment was created with a plastic box homogeneously wrapped in white paper.

Hypocotyl width and epidermal cell length was obtained from hypocotyl SEM images (see above) from 3 dps seedlings grown in normal light conditions. SEM images allowed the hypocotyl files of large non-dividing cells to be distinguished from files of small cells. In Arabidopsis hypocotyls, the large non-dividing cells are positioned outside of a single cortical cell file, and are characterized by the expression of *GL2*, the marker for root non-hair cells (Daher et al. 2018; Gao et al. 2022). Hypocotyl width measurements were made at the rootward, center, and shootward regions of the hypocotyl and then values were aggregated for the quantification shown.

### Chemical Treatments

Treatments with small molecules were performed using *pWFL:WFL-GFP* in the Col-0 background seedlings grown on 0.5X MS plates grown for 5 dps. Seedlings were then incubated in liquid 0.5x MS containing one or a combination of the following chemicals: Brefeldin A (BFA, Sigma-Aldrich) was dissolved in dimethyl sulfoxide (DMSO, Calbiochem, Cat #317275) in 50 mM stocks and added to the media at a final concentration of 50 µM for 1 hour or other indicated times. Cycloheximide (CHX, Sigma-Aldrich, Cat #C7698) was dissolved in DMSO and added from a 50 mM aqueous stock to a final concentration of 50 µM for 2 hours or other indicated time. Wortmannin (Wm, Sigma-Aldrich) was dissolved in DMSO and used at 33 µM for 2 hours. In control (mock) experiments, seedlings were incubated in the same media containing an equal amount (0.05% to 0.1%) of the solvent. Washout was performed by transferring seedlings to new 0.5x MS media containing one or more of the treatments as described in the results section.

### qRT-PCR Analysis

RNA was isolated with Qiagen’s RNeasy Plant Mini Kit from whole seedlings at 7 dps from three independent biological replicates. First-strand cDNA was synthesized from 1 μg total RNA with RevertAid First Strand cDNA Synthesis and the oligo(dT)_18_ primer (Thermo Scientific). qRT-PCR reactions were set up in technical replicates using IQ SYBR Green Supermix (BioRad). Cycling conditions: 95°C for 3 min followed by 40 cycles of 95°C for 10s and 57°C for 20 s. Standard curves were performed at least in duplicate. Primer efficiencies were measured as 90-110% for all primers. See Table S1 for primer sequences. Transcript levels were calculated using the software CFX Manager (Bio-Rad), normalizing to *PP2A/*AT1G13320 (Czechowski et al. 2005).

### Statistical analysis

All pairwise comparisons were performed with GraphPad Prism 10 (www.graphpad.com) or ggplot2 (https://ggplot2.tidyverse.org) package in RStudio, for parametric (*t*-test) and non-parametric analysis (Wilcoxon). Multiple comparison one-way ANOVA with Dunn’s multiple comparisons post-test was used in GraphPad Prism 10 (www.graphpad.com).

## RESOURCE AVAILABILITY

The WALLFLOWER (WFL) accession number is At5g24100 and the WALLFLOWER-RELATED 1 (WFLR1) accession number is At5g53320. The data and plant and molecular materials underlying this article will be shared upon reasonable request to the corresponding author.

### Lead Contact

Requests for further information and resources should be directed to and will be fulfilled by the lead contact, Jaimie M. Van Norman.

## Materials Availability

The plant and molecular reagents underlying this article will be shared without restriction upon reasonable request to the lead contact.

## Data and Code Availability

All data reported in this paper will be shared by the lead contact upon request. This paper does not report original code. Any additional information required to reanalyze the data reported in this paper is available from the lead contact upon request.

## ACKNOWLEDGMENTS

We thank Roya Campos, Jason Goff, Carolyn Rasmussen and Patricia Springer (UCR) for helpful discussions during the early phases of this work. We are also very grateful to Michael Guzman (UCR) for agreeing to include *wfl* in one of his ongoing hypocotyl elongation assays. We thank Siobhan Braybook (UCLA) for helpful guidance using MorphGraphX, sharing PPP-CONT cantilevers and advice on the model to be used for data processing. We thank Jeff Long (UCLA) for providing access to the vibratome. We acknowledge the use of FLIM confocal microscopy at the Advanced Light Microscopy/Spectroscopy Laboratory (RRID: SCR_022789) and at the Leica Microsystems Center of Excellence, and also the use of NanoWizard® 4a BioScience Atomic Force Microscope at the Nano and Pico Characterization Lab (RRID:SCR_022924) at the California NanoSystems Institute at UCLA. We appreciate access to and assistance at the IIGB Genomics Core and Microscopy Core (UCR). We also thank Erin Sparks (Delaware Biotechnology Institute) for providing the NorfT version of the dpGreen Gateway compatible destination vector and Sebastien Santini (CNRS/AMU IGS UMR7256) and the PACA Bioinfo platform for the availability and management of the phylogeny.fr website. The initial phases of this work was supported by Initial Complement (IC) funds from the University of California, Riverside, USDA-NIFA-CA-R 587 BPS-5156-H, NSF CAREER award #1751385 to J.M.V.N., and NSF-GRFP award #DGE 588 1326120 to J.N.T. Support for the later phases, including phenotypic analyses, 3D imaging, and cell wall imaging and analyses came from Initial Complement (IC) funds from the University of California, Los Angeles to J.M.V.N. A.A.B. acknowledges National Science Foundation (NSF) support via Major Research Instrument (MRI) DMR Project #2019056 entitled “Development of a Cryogenic Integrated Micro-Raman-Brillouin-Mandelstam Spectrometer.”

## AUTHOR CONTRIBUTIONS

Initial stages of the work at UC Riverside: Conceptualization: J.M.V.N.; Methodology and Investigation: J.M.V.N., J.N.T., and C.R-F.; Resources: J.M.V.N. and J.N.T.; Writing - first manuscript (posted in BioRxiv): J.N.T., C.R-F., and J.M.V.N.; Visualization: J.N.T. and C.R-F.; Supervision: J.M.V.N.; Funding Acquisition: J.M.V.N. and J.N.T.

Later stages of work at UC Riverside and UC Los Angeles: Conceptualization: P.P-H. and J.M.V.N.; 3D imaging: P.P-H.; *wfl* growth phenotypic analysis: P.P-H, R.E.D.; Brillouin spectroscopy measurements and analysis: D.W., P.P-H and A.A.B, FLIM measurements and analysis, PP-H and H.M, AFM measurements, P.P-H, Double mutant analysis *wfl* and related LRR-RLKs: R.E.D.; Visualization: P.P-H., R.E.D.; Writing - updated manuscript: P.P-H; Review and editing: J.M.V.N., P.P-H., R.E.D., C.R-F., and J.N.T.; Funding Acquisition: J.M.V.N.

## SUPPLEMENTAL INFORMATION

Figures S1-S5, Table S1, Supplementary videos 1-2

---

**Figure S1.**
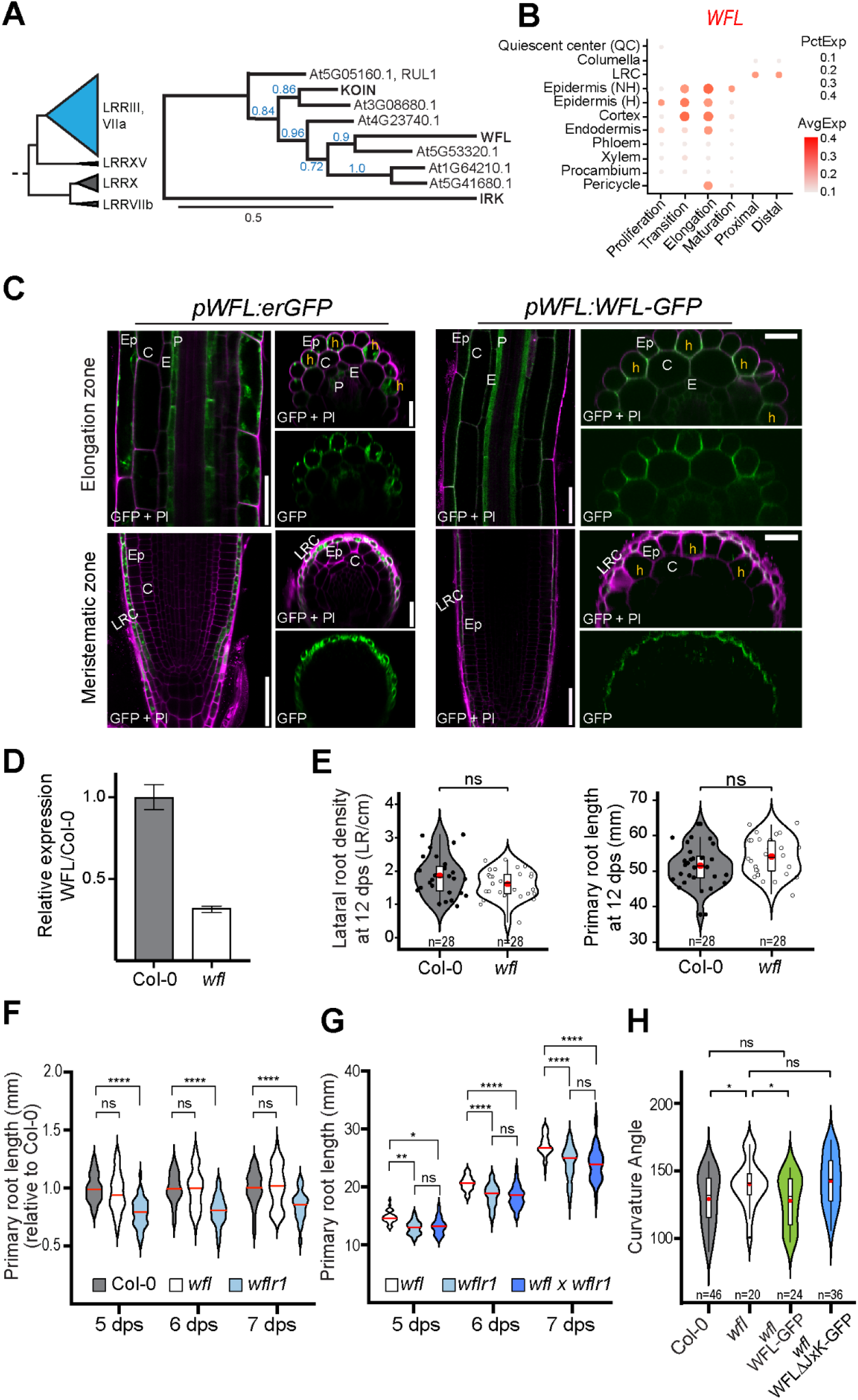
*WFL* promoter activity and protein accumulation and various phenotypic assays. **A**) Left, Partial phylogenetic tree adapted from (Man et al. 2020) with only Arabidopsis proteins of interest included. Triangles drawn to scale to represent the number of proteins included: LRRIII, VIIa - 54; LRRXV - 2, LRRX - 11, and LRRVIIb - 3. Subgroup LRRIII, VIIa (blue) contains WFL and several other Arabidopsis LRR-RLKs with polar accumulation in root cell types, including KOIN (Rodriguez-Furlan et al. 2022) and IRK (Campos et al. 2020). *Right*, Unrooted phylogram with bootstrap support values in blue. Members of the LRRIII/VIIa subgroup, including KOIN (At5G53800) and WFL (At5G24100) from LRRIII, with IRK (At3G56370) included as a more distantly related subfamily member (representing subgroup VIIa). Legend shows branch length 0.5. Phylogram built at https://www.phylogeny.fr/alacarte.cgi using “a la carte mode” (Dereeper et al. 2008). **B**) *WFL* mRNA expression data from atlas ARVEX - https://shiny.mdc-berlin.de/ARVEX (Shahan et al. 2022). Color intensity represents *WFL* expression level (AvgExp), and dot size depicts the percentage of cells with that expression (PctExp). C) Roots stained with propidium iodide (PI, magenta) expressing (*Left panels*) *pWFL:erGFP* and (*Right panels*) *pWFL:WFL-GFP*. Adjacent panels show GFP alone (Green) or GFP + PI signal in the meristematic and elongation zones. **D**) RT-qPCR shows reduced *WFL* transcript levels with 3 technical replicates conducted per biological replicate (of 3) with similar results obtained in all. Plot shows mean ± SEM. E) Emerged lateral root density (LR number / primary root length) and corresponding primary root length measured in Col-0 and *wfl* seedlings at 12 dps. N = 28 roots **F**) Primary root length of *wfl* and *wflr-1* (At5G53320) roots relative to Col-0 at 5, 6, and 7 days post-stratification (dps). Violin plot shows the data and distribution of all data points relative to the average of Col-0 at each day. Median is a red horizontal line. One-way ANOVA with Dunn’s multiple comparisons test. Col-0: 5 dps n = 81, 6 dps n = 66, 7dps n = 51; *wfl*: 5 dps n = 70, 6 dps n = 58, 7 dps n = 44;*wfl*1: 5-7 dps n = 34. **** (p-value < 0.0001). G) Primary root length wfl, wflr1, and *wfl wflr1* double mutants at 5, 6, 7 dps. Violin plot shows data distribution and the median value as a red horizontal line. One-way ANOVA with Dunn’s multiple comparisons test. *wfl*: 5 dps n = 68, 6 dps n = 54, 7 dps n = 44; *wflr1*: 5 dps n = 59, 6 dps n = 53, 7 dps n = 43; *wfl wflr1*: 5 dps n = 64, 6 dps n = 53, 7 dps n = 41. ns = no significance (p-value > 0.05), * (p-value = 0.0335), ** (p-value = 0.0033), **** (p-value < 0.0001). H) Angle of the root curvature during a gravitropic reorientation (180°). Violin plots show data distribution with box plots indicating the first and third quartiles and black dots showing outliers. ‘n’ indicates the number of roots examined. The median and average values are shown by a line and red dot, respectively. Unpaired two-tailed t-test with ns = no significance (p-value > 0.05) and * = p-value <0.05. Supplementary Figure S1 supports data presented in Figure 1.

**Figure S2.**
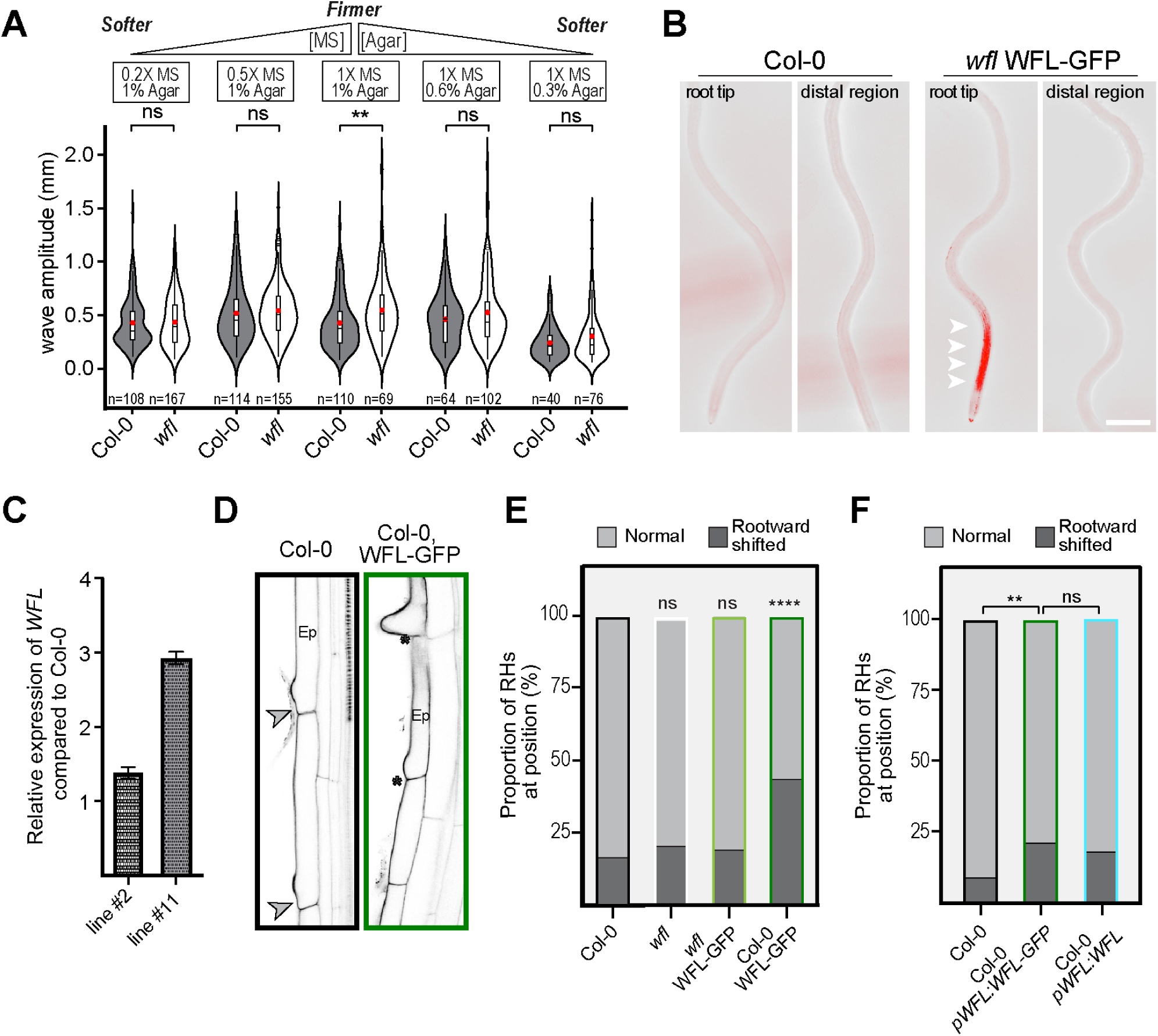
WFL-GFP accumulates strongly in the elongation zone and root hair positioning is altered when extra copies of WFL are present. A) Quantification of wave amplitude for Col-0 and *wfl* seedlings grown on varying amounts of mineral salt levels (MS) and gelling agent (agar). Violin plots show data distribution with box plots indicating the first and third quartiles and dots showing outliers. ‘n’ indicates the number of waves examined. The median and average values are shown by a line and a red dot, respectively. Unpaired two-tailed *t*-test with ns = no significance (p-value > 0.05), ** (p-value < 0.01). B) Wild type (Col-0) and *wfl pWFL::WFL-GFP* seedlings at 5 dps grown on inclined (20°) plates and imaged with fluorescence microscopy (GFP displayed in red). White arrowheads indicate strong WFL-GFP signal at the elongation zone. Scale bar = 0.5 mm. C) *WFL* transcript levels in two independent lines expressing extra copies of *WFL* (Col-0, *pWFL::WFL-GFP*). *WFL* relative expression was measured by RT-qPCR relative to the expression in untransformed Col-0. Plot shows mean ± SEM. D) Confocal images of root hair cells in Col-0 and Col-0 *pWFL:WFL:GFP*. Gray arrowhead indicates normal gap between the root hair and the rootward end of the cell. Asterisk indicates absence of a gap. E) Proportion of root hairs classified as normally positioned (light grey) or shifted downwards (dark grey) in different genetic backgrounds (n= 15 roots). *t*-test, ns = not significant (p > 0.05), ** p < 0.01, and **** p < 0.0001 . F) Proportion of root hairs classified as normally positioned (light grey) or shifted downwards (dark grey) in Col-0 and Col-0 expressing WFL with and without GFP. Supplementary Figure S2 supports data presented in Figure 1 and 2.

**Figure S3.**
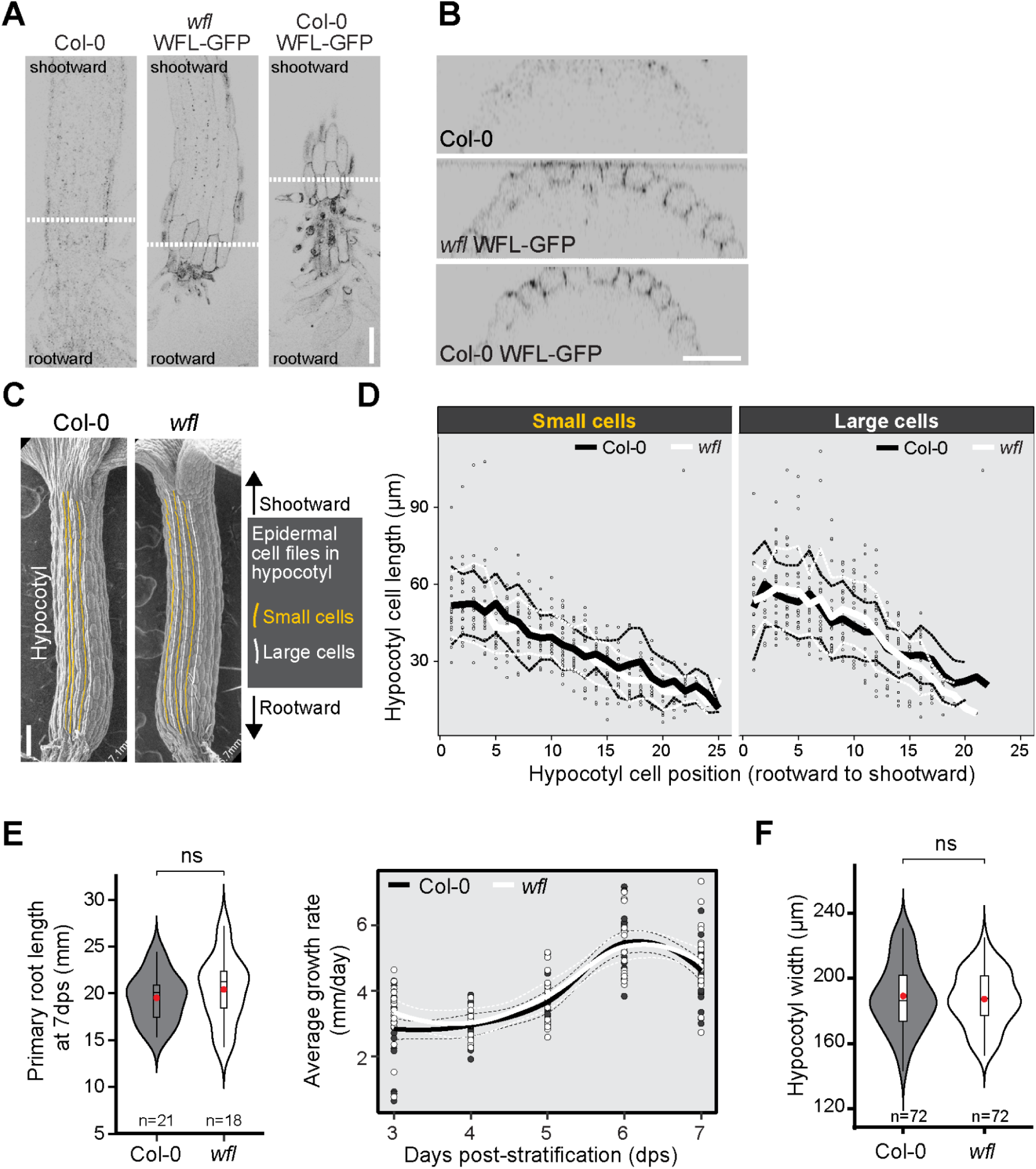
Loss of *WFL* does not affect hypocotyl width or epidermal cell length in normal light conditions. **A**, **B)** Confocal images of WFL-GFP in Col-0 and *wfl* seedlings at 2.5 dps. **A)** Longitudinal optical sections (*xy* planes) at the surface of hypocotyls. Black arrowhead indicates root-hypocotyl junction and white dashed line indicates position of transverse optical sections in B. Scale bar = 60 µm. **B)** Transverse optical sections (*xz* planes) at positions in A. Scale bar = 60 µm. **C)** Scanning electron microscopy (SEM) images from Col-0 and *wfl* hypocotyls at 3 dps allows for discrimination between cells that are small (yellow vertical lines) and large (white vertical lines) and measurement of their lengths. Scale bar = 100 µm. **D)** Scatter plot of hypocotyl epidermal cell length arranged by their position (rootward to shootward). Graph shows individual data points with mean value (thick line) and standard deviation (dashed lines). n= 886 and 922 cells for Col-0 and *wfl*, respectively. **E)** Total root length and root growth rate from seedlings grown on media without sucrose (same growth conditions for hypocotyl assays). *Left panel*, Total primary root length at 7 dps. Violin plot shows data distribution with the box plots indicating the first and third quartiles and median and average values shown with a line and red dot, respectively. Unpaired two-tailed *t*-test. No statistically significant difference (ns, p-value > 0.05). *Right panel*, Corresponding root growth rates. Graph shows individual data points with mean value (thick line) and standard deviation (dashed lines). **F)** Hypocotyl width of Col-0 and *wfl* seedlings at 3 dps. The violin plot shows data distribution with the box plot indicating the first and third quartiles, the median value shown by a line, and the average by a red Unpaired two-tailed *t*-test with Welch’s correction. No statistically significant difference (ns, p-value > 0.05). Supplementary Figure S3 supports data presented in Figure 3.

**Figure S4.**
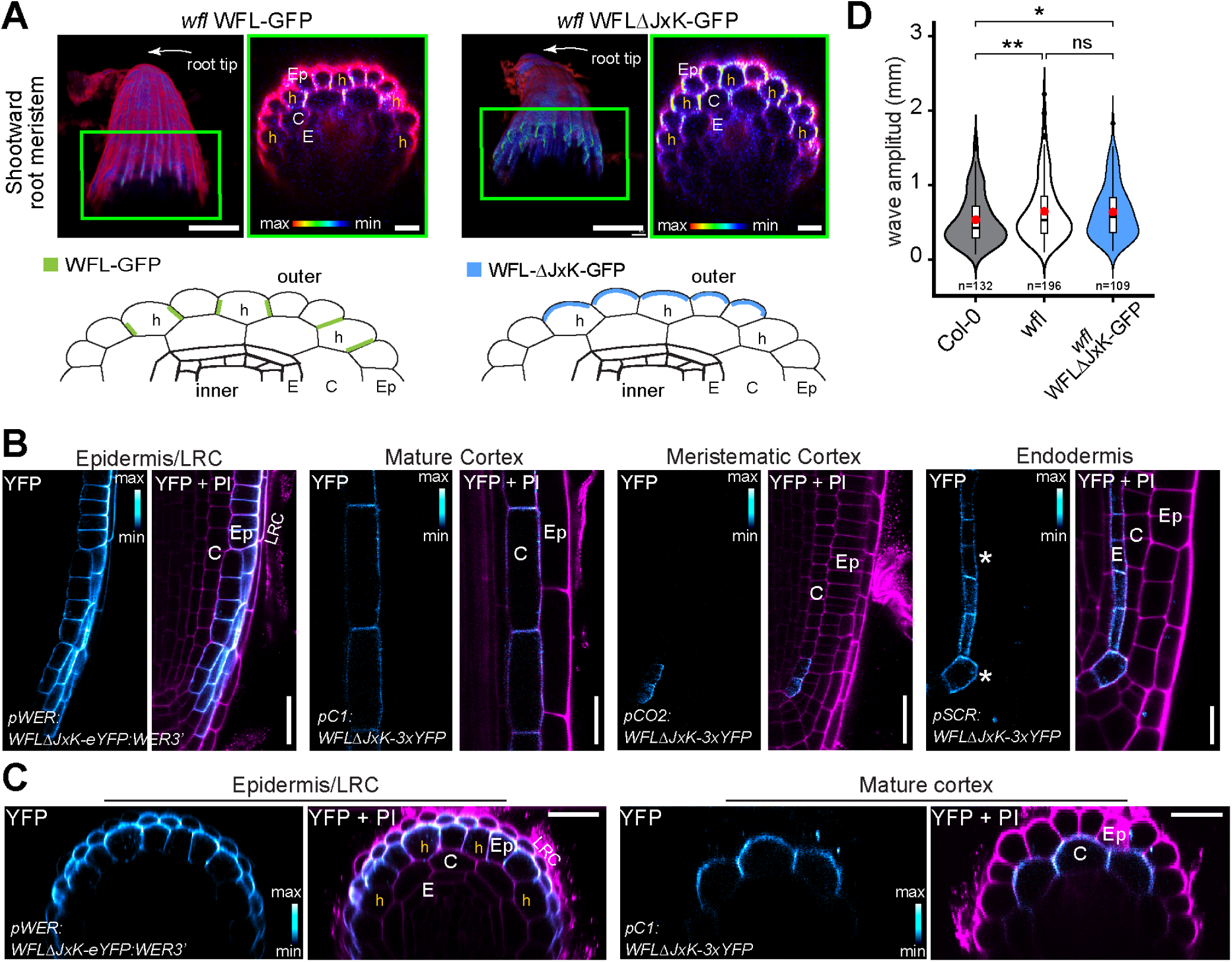
WFLΔJxK-GFP predominantly localizes to the outer polar domain of root cell types. **A)** *Upper panels*, Confocal images of the shootward portion of root meristems stained with propidium iodide (PI, magenta). *Left*, 3D projections and (right) *xz* planes for each genotype. WFL-GFP accumulates at the circumferential polar domain of hair cells (as seen in Figure 1b), whereas WFLΔJxK-GFP accumulates at the circumferential and outer polar domains of all epidermal cells. *Lower panels*, Schematic representations of WFL-GFP (green) and WFLΔJxK-GFP (blue) accumulation in the root epidermal cells in the shootward portion of the meristematic zone. Scale bar = 50 µm and 10 µm for left and right images, respectively. **B, C)** Roots expressing WFLΔJxK-eYFP/3xYFP driven by cell layer-specific promoters and stained with propidium iodide (PI, magenta). Adjacent panels show YFP alone (shown with signal intensity color scale) and YFP + PI merged in (**B**) median longitudinal planes and (**C**) transverse planes. In the ground tissue initials and endodermis of lateral roots (far right panels (**B**)), WFLΔJxK-eYFP displays nonpolar accumulation (white asterisks). Scale bars: 25 μm. **D)** Quantification of wave amplitude for Col-0, *wfl*, and *wfl* WFLΔJxK-GFP. Violin plots show data distribution with box plots indicating the first and third quartiles and dots showing outliers. ‘n’ indicates the number of roots examined. The median and average values are shown by a black line and red dot, respectively. Unpaired two-tailed *t*-test was used. ns = no significance (p-value > 0.05), * (p-value < 0.05), ** (p-value < 0.01). Supplementary Figure S4 supports data presented in Figure 5.

**Figure S5.**
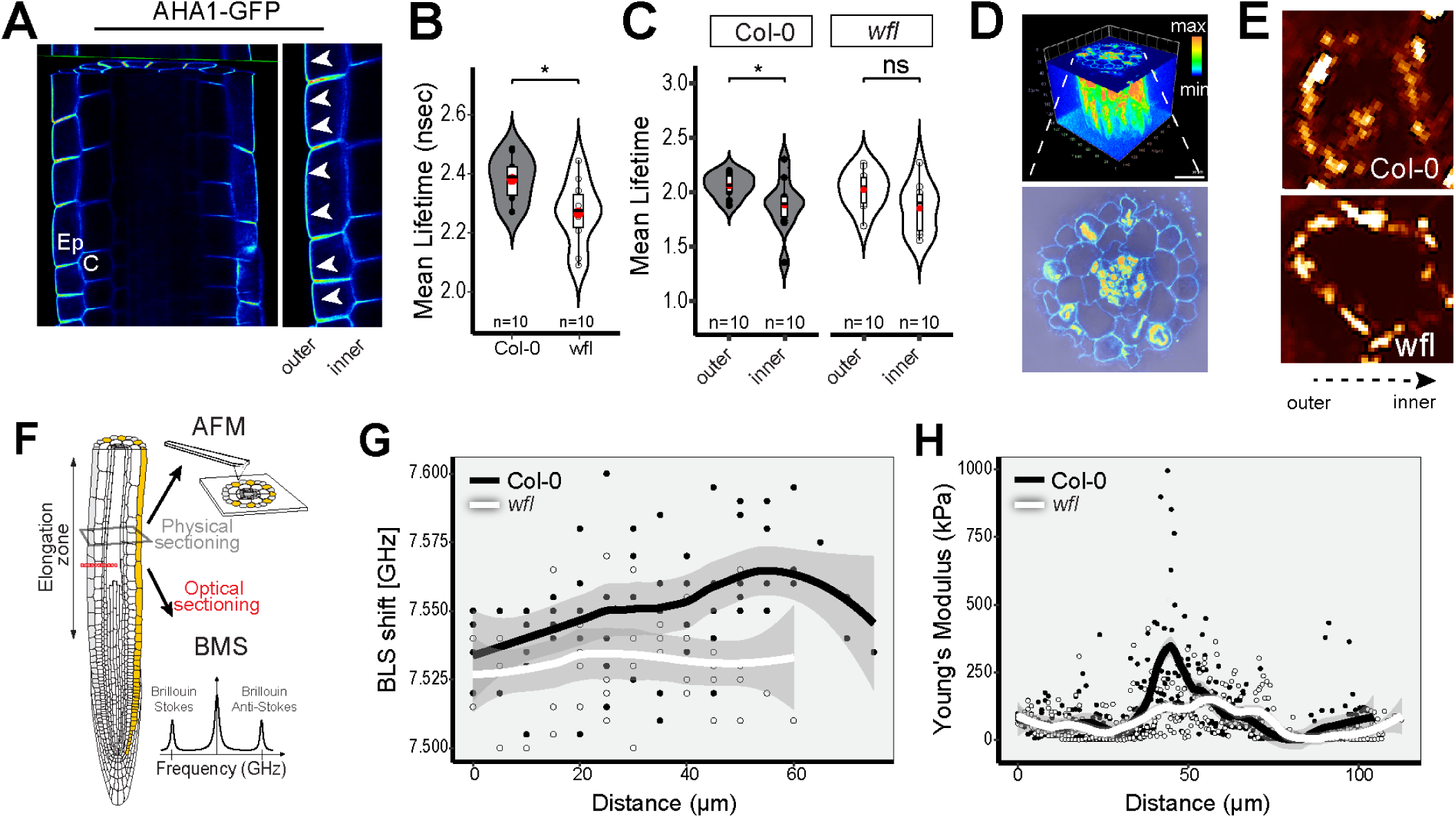
Lateral cell wall properties in elongating *wfl* epidermal cells are different from wild type. **A)** 3D confocal imaging of roots expressing a *pAHA1::AHA1-GFP* reporter and close up of an epidermal (Ep) cell file. Arrowheads point to localization of the plasma membrane H+-ATPase AHA1-GFP to the outer polar domain of epidermal cells as they begin to elongate. **B)** Mean fluorescence lifetime obtained from optical cross-sections at the elongation zone using aFLIM. nsec = nanoseconds. Violin plots show data distribution with box plots indicating the first and third quartiles. Median and average values are indicated with a line and a red dot, respectively. Unpaired two-tailed *t*-test. ns = no significance (*p*-value > 0.05) and * (*p*-value < 0.05). n = number of cells, from 10 roots per genotype. **C)** Mean lifetime obtained from ROIs at the outer and inner sides of the epidermal cells from FLIM images. Violin plots show data distribution with box plots indicating the first and third quartiles with black dots showing outliers. Median and average values are indicated with a line and a red dot, respectively. Unpaired two-tailed *t*-test. ns = no significance (*p*-value > 0.05) and * (*p*-value < 0.05). n = number of cells, from 11 roots per genotype. **D)** Test of the root integrity for AFM experiments after physical sectioning. *Top*, 3D confocal imaging of an agar-embedded root after sectioning with a vibratome. Sections are 200 μm thick. Cell walls are stained with Propidium Iodide (PI, blue to orange color scale). *Bottom*, confocal image of the cross-section at the top of the agar section. PI channel (blue to orange) and brightfield channel are merged. **E)** Map of Young modulus values of single epidermal cells obtained by AFM in advanced quantitative imaging mode. **F)** Root schematic displaying the physical sectioning for Atomic Force Microscopy (AFM) and optical sectioning for Brillouin–Mandelstam light scattering (BMS) microscopy. **G)** Brillouin shift values along the radius of the root from the outside edge towards the center. Continuous lines indicate a polynomial fitting and gray area around each line indicates a 95% confidence interval. n = 5 roots per genotype. **H)**, Young modulus values obtained by AFM across the root diameter. Values obtained from a line profile crossing the entire root cross section connecting the 8 cardinal points in that plane. Continuous lines indicate a polynomial fitting and the gray area around each line indicates the 95% confidence interval. n = 4 line profiles per root, from 3 roots per genotype. Supplementary Figure S5 supports data presented in Figure 6.

**Table S1.**
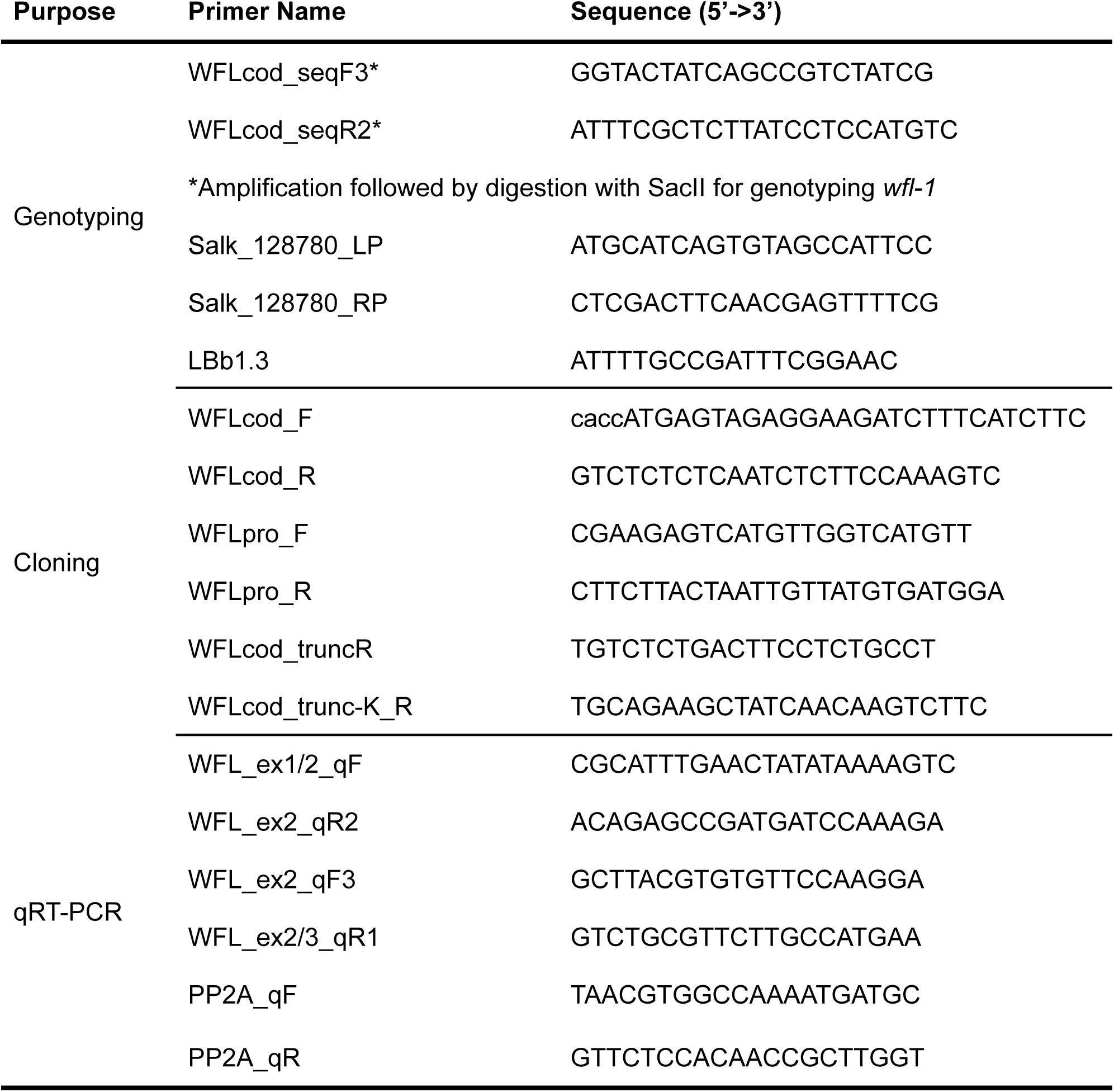
List of primers used in this study for genotyping and cloning.

